# In vivo screening identifies SLC38A3 as a metabolic booster for CAR T cell therapy

**DOI:** 10.64898/2026.11.24.768308

**Authors:** Wentao Li, Gernot Wolf, Jenny Lin, Nicole Pranckevicius, Rita Simões-Faria, Raphael Bednarsky, Mustapha Jaiteh, Cecilia Georges, Philipp Muench, Francesco Piras, Sabrina Ladstätter, Thomas Krausgruber, Laura P.M.H. de Rooij, Giulio Superti-Furga, Christoph Bock

**Author notes:** Correspondence (C.B.).

## Abstract

Chimeric antigen receptor (CAR) T cells are highly effective against certain blood cancers, but struggle in the hostile tumor microenvironment (TME) of solid tumors. We developed the CARGO method for in vivo gain-of-function screening in CAR T cells and used it to equip CAR T cells with enhanced access to nutrients inside the TME, by overexpressing 452 solute carrier (SLC) transmembrane transporters. We identified SLC38A3, a sodium-coupled amino acid transporter not endogenously expressed in T cells, as an effective booster of CAR T cell performance in xenograft models of hepatocellular carcinoma (Huh7, HepG2) and ovarian adenocarcinoma (SKOV3). Ectopic expression of SLC38A3 in CAR T cells enhanced their glutamine uptake, improved metabolic fitness, and promoted a stem-like memory phenotype, which resulted in superior antitumor efficacy and durable protection upon tumor rechallenge in glutamine-restricted solid tumors. In summary, our study shows that equipping CAR T cells with non-native nutrient transporters can confer enhanced metabolic resilience, and it establishes the feasibility and utility of in vivo open reading frame (ORF) screens in CAR T cells to tackle the hostile TME of solid tumors.

**Highlights:**

- In vivo gain-of-function screening with 452 solute carrier (SLC) transmembrane transporters to find metabolic boosters of CAR T cell function
- Discovery and validation of improved in vivo performance of CAR T cells that overexpress the amino acid transporter SLC38A3 in xenograft tumors
- SLC38A3 enhances the glutamine uptake of CAR T cells, resulting in improved persistence and metabolic function in nutrient-restricted solid tumors
- CAR T cells expressing SLC38A3 show a stem-like T cell phenotype and achieve superior efficacy in additional models of liver and ovarian cancer

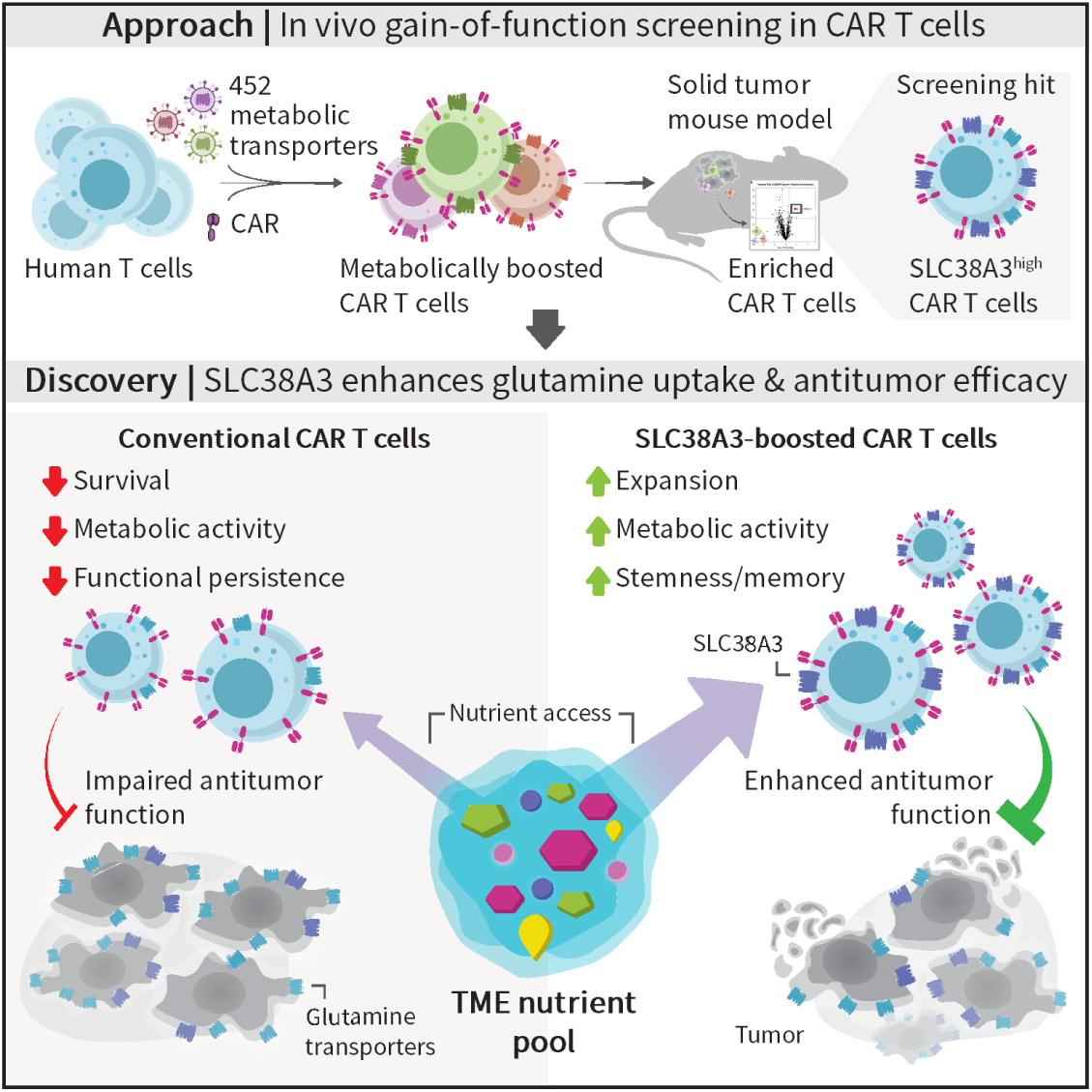

## Introduction

Chimeric antigen receptor (CAR) T cells constitute a medical breakthrough for the treatment of B cell malignancies^1,2^. However, the translation of this success to patients with solid cancers has been challenging. In solid tumors, metabolic processes create a hostile tumor microenvironment (TME)^3,4^ that is often characterized by nutrient deprivation and accumulation of oncometabolites^5,6^. For example, glutamine-addicted cancer cells have the ability to outcompete immune cells in the uptake of this amino acid^7,8^. As a consequence, tumor-infiltrating lymphocytes, including CAR T cells, can undergo metabolic exhaustion and lose their antitumor efficacy^9–11^. Because mitochondrial fitness is important for generating long-lived, stem-like memory T cells, metabolic constraints in the TME not only compromise immediate tumor killing but also undermine the durability of antitumor responses.

Several approaches have been developed for enhancing the metabolic fitness of CAR T cells in solid tumors^12,13^. For example, knockout of the adenosine A2A receptor (A2AR) in CAR T cells improved their antitumor function by reducing adenosine signaling^14^; knockout of methylation-controlled J protein (MCJ), a negative regulator of mitochondrial respiration, elevated mitochondrial metabolism and antitumor activity in CAR T cells^15^; and knock-in of the mitochondrial enzyme proline dehydrogenase 2 (PRODH2) increased CAR T cell-mediated killing^16^. These approaches have focused on the tuning of metabolic genes and pathways that are active in T cells, while the integration of genuinely non-native metabolic capabilities into CAR T cells has yet received little attention.

Most metabolites cannot freely diffuse across the plasma membrane but require specialized transporters for cellular uptake^17,18^, with solute carrier (SLC) proteins playing a central role^19^. Tumor cells can enhance their access to nutrients by upregulating SLCs^20–22^, suggesting that ectopic expression of SLC genes among CAR T cells could help them compete for scarce metabolites. Previous studies focused on endogenous T cell candidates and found that increased expression of the glucose transporter GLUT1 (SLC2A1) and the amino acid transporters SLC7A5 and SLC7A11 can improve certain aspects of CAR T cell function^23–25^. However, these hypothesis-driven efforts evaluated only very few SLC genes, leaving the wider potential of metabolic enhancement in CAR T cells untested.

In this study, we sought to enhance the metabolic fitness of CAR T cells in solid tumors by improving their access to nutrients through overexpression of SLC transmembrane transporters. We developed the CARGO (“CAR + Gene Overexpression”) method for gain-of-function screens with a single lentiviral vector that co-delivers the CAR together with an open reading frame (ORF) library comprising 452 codon-optimized SLC cDNAs^26^, to individually boost the expression levels of essentially all SLCs in human primary CAR T cells. We opted for ORF overexpression over CRISPR activation (CRISPRa) as CRISPRa struggles to activate genes that are transcriptionally silent^27,28^ and to further upregulate genes that are already highly expressed at baseline^29^. Thus, our screening method is not constrained by endogenous gene expression in T cells – it allows us to introduce non-native uptake capabilities and thereby expands the metabolic repertoire that CAR T cells can have access to. Moreover, the CARGO method is fully generalizable to ORF libraries comprising other proteins with potential benefit to CAR T cells.

We initially pursued a series of in vitro screens to enhance the metabolic fitness of CAR T cells, but their results did not translate to preclinical in vivo models (**Supplementary Note**). Consistent with recent reports^30,31^, this finding highlights the limitations of in vitro models for studying CAR T cell fitness, and prompted us to pursue further gain-of-function screens exclusively in vivo, in xenograft tumors that recapitulate the metabolic challenges that CAR T cells encounter in solid tumors. Our screening established a comprehensive assessment of SLCs as metabolic enhancers in CAR T cells and identified ectopic expression of SLC38A3, an amino acid transporter not endogenously expressed in T cells, as a booster of CAR T cell performance through enhanced nutrient access in the TME. We extensively tested the functional effects conferred by SLC38A3 and found that it promotes a stem-like CAR T cell phenotype and establishes durable antitumor efficacy in glutamine-restricted solid tumors. Our study thus demonstrates the power of gain-of-function in vivo screening for metabolic boosters of CAR T cell function.

## Results

### CARGO supports in vivo gain-of-function screening in CAR T cells

To screen for metabolic boosters of CAR T cell function in xenograft mouse models of solid tumors, we developed and validated a single-vector system for co-delivery of a CAR and a custom ORF library (**Fig. 1a** and **Fig. S1**). Our CARGO vector includes key features for efficient in vivo screens, such as effective quantification by barcode sequencing and built-in support for clonality assessment and correction with unique molecular identifiers (UMIs).

**Figure 1.**
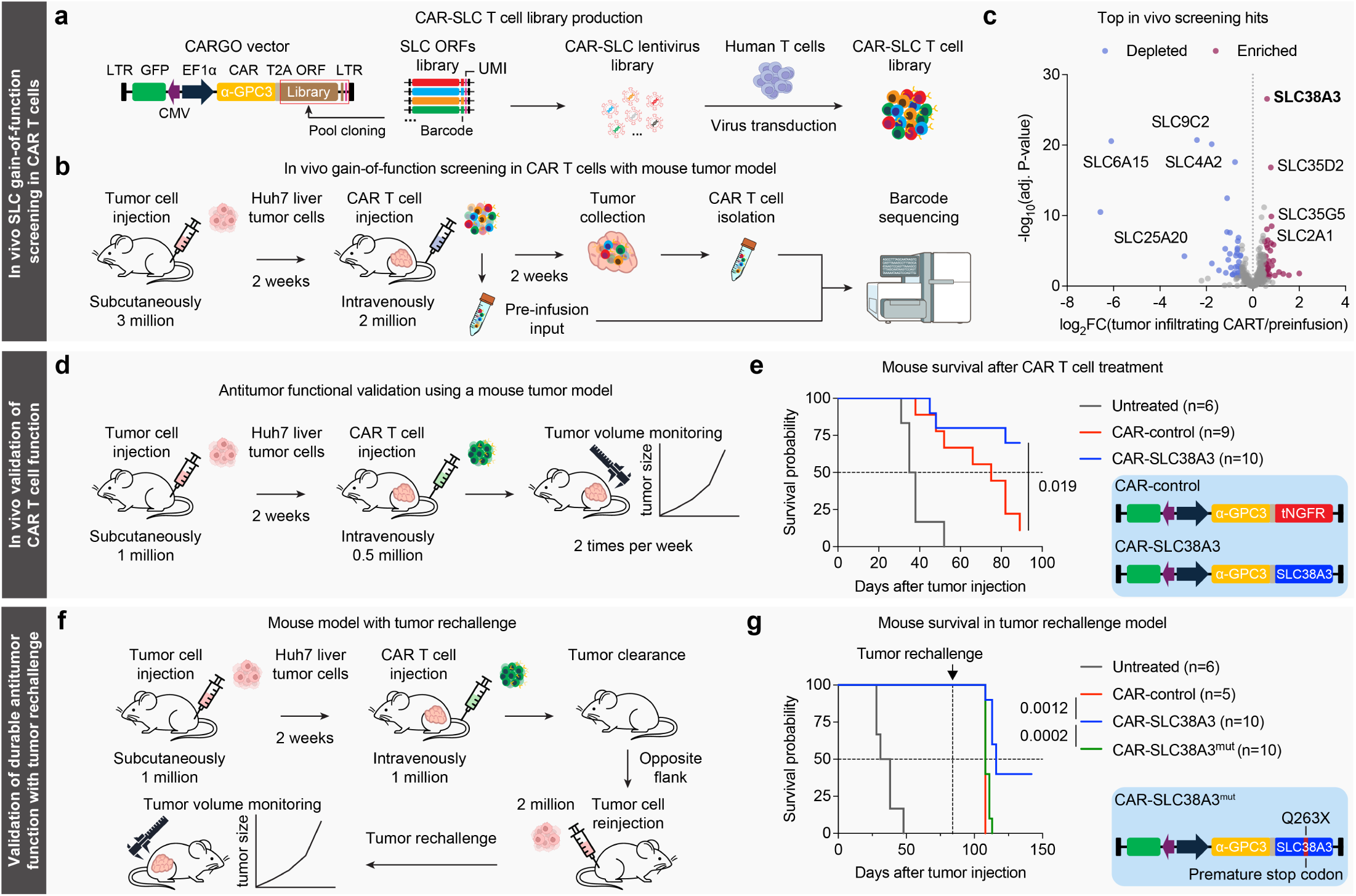
In vivo gain-of-function screening identifies SLC38A3 as a metabolic booster of CAR T cell function. (a) Schematic of the CARGO vector for in vivo screening and of the CAR-SLC T cell library. 452 codon-optimized solute carrier (SLC) cDNAs were cloned in bulk into the CARGO vector to produce lentivirus for gain-of-function screens. Human T cells were then transduced with the lentivirus, resulting in a ready-to-screen CAR T cell pool. (b) Outline of the in vivo gain-of-function screening with human CAR T cells targeting xenograft liver tumors. (c) Volcano plot showing changes in SLC abundance among tumor-infiltrating CAR T cells compared to the pre-infusion cells. Purple dots indicate significantly enriched SLCs (log_2_FC > 0.5, FDR-adjusted *p*-value < 0.05), blue dots indicate significantly depleted SLCs (log_2_FC < -0.5, FDR-adjusted *p*-value < 0.05), and grey dots indicate unchanged SLCs. Screens were performed independently for three T cell donors with three to six mice each. (d) Outline of in vivo validations for SLC38A3-expressing CAR T cells in xenograft liver tumors (Huh7). (e) Survival curves for tumor-bearing mice left untreated (grey), injected with control CAR T cells (which express truncated tNGFR^23,67–69^, red), or with SLC38A3-expressing CAR T cells (blue). The CAR delivery vectors are shown on the right. Experiments were performed independently for two T cell donors with five mice each. (f) Outline of the tumor rechallenge model for assessing persistence and long-term function of the CAR T cells. Mice that had cleared the primary tumor were rechallenged on day 84 by injecting Huh7 tumor cells in the opposite flank without giving a new dose of CAR T cells, thus relying on persistent CAR T cells from the initial dose. (g) Survival curves for the tumor rechallenge model, showing mice left untreated (grey) or previously injected with control CAR T cells (red), CAR T cells expressing SLC38A3 (blue), or CAR T cells expressing the transport-deficient SLC38A3^mut^ variant (green). Experiments were performed independently for two T cell donors with five mice each. *P*-values in (e) and (g) were determined by the log-rank (Mantel-Cox) test.

We equipped the CARGO vector with an anti-GPC3 CAR for in vivo screens in Huh7 liver tumors, a xenograft model that reflects the metabolic challenges of hepatocellular carcinoma. We then cloned our ORF library comprising 452 codon-optimized SLC cDNAs (**Fig. S1a and Table S1**) into this vector, prepared lentivirus, transduced primary human T cells, and expanded these CAR T cells, ready to be injected into Huh7 tumor-bearing mice (**Fig. 1b**).

Based on a series of titration experiments, we selected the number of tumor cells and the corresponding number of CAR T cells such that the CAR T cells achieved effective in vivo tumor control while preserving sufficient tumor mass that we could harvest for extracting tumor-infiltrating CAR T cells as our screening readout (**Fig. S1b,c**).

We established the feasibility of in vivo screening using this setup by isolating CAR T cells from the tumors, performing ORF barcode sequencing, and comparing them with pre-infusion CAR T cells. We confirmed the presence of almost all SLCs in the plasmid library (99.3%) and recovered them from the CAR T cells at the start (98.8%) and end (98.6%) of the in vivo screen (**Fig. S1d; Table S2**). We retained robust barcode sequencing coverage across the ORF size range from 327 to 4077 base pairs (**Fig. S1e**), backed by a large number of independent clones (assessed by UMI counting) for all but the most depleted genes across three T cell donors (**Fig. S1f; Table S3**).

In summary, we established a method for co-delivering a CAR and an ORF library into human primary T cells and for screening the modified CAR T cells in terms of their ability to infiltrate, survive, and proliferate in solid tumors.

### Overexpression screening identifies SLC38A3 as a booster of CAR T cell function

Following subcutaneous injection of the tumor cells and intravenous injection of the engineered CAR T cells two weeks later, we harvested the tumors after another two weeks, isolated tumor-infiltrating CAR T cells, and quantified the enrichment or depletion of CAR T cells expressing each SLC based on barcode sequencing and comparison to pre-infusion CAR T cells. We identified multiple SLCs that were significantly enriched or depleted in our screen (**Fig. 1c and S1g; Table S2**), suggesting that their overexpression fosters (for enriched SLCs) or impairs (for depleted SLCs) the CAR T cells’ ability to establish and maintain a strong presence inside these tumors.

For example, SLC4A2 and SLC9C2 were significantly depleted in our screen, possibly because their overexpression interferes with ion regulation and pH balance^32^. Conversely, several SLC35 family members^33^ were significantly enriched, indicating that their expression is beneficial for the CAR T cells’ ability to infiltrate and persist in these tumors. We also rediscovered GLUT1 (SLC2A1), the previously identified transporter^23,24^ that motivated this study, as a highly significant screening hit (FDR-adjusted *p*-value: 3.0×10^-7^), supporting the validity of our screen.

The most significant and arguably the most interesting hit from our screen was SLC38A3 (FDR-adjusted *p*-value: 2.8×10^-27^). SLC38A3 is a sodium-coupled amino acid transporter that mediates the uptake of glutamine, asparagine, and histidine^34–36^. It is not expressed in T cells but highly expressed in hepatocellular carcinoma tumor cells^37^ (**Fig. S1h**), suggesting that this transporter confers tumor cells with enhanced access to key amino acids, and that its ectopic expression may restore the CAR T cells’ ability to compete with the tumor cells for nutrient access.

In summary, we conducted in vivo gain-of-function screening in CAR T cells that infiltrate xenograft liver tumors, which identified an SLC that is not endogenously expressed in T cells as a putative booster of CAR T cell function.

### SLC38A3 expression robustly enhances CAR T cell efficacy in vivo

To validate the beneficial effect of SLC38A3 on the CAR T cells’ antitumor efficacy, we compared constitutively SLC38A3-expressing CAR T cells with matched control CAR T cells in our Huh7 tumor model (**Fig. 1d and S2a**). The SLC38A3-expressing CAR T cells achieved superior tumor control and significantly extended survival (*p*-value: 0.019), with 70% of the mice surviving beyond 90 days compared to 11% in the control group (**Fig. 1e and S2b**).

To confirm that this effect was driven by the transporter function of SLC38A3, we tested a previously reported transport-deficient mutant^38^ and found that CAR T cells expressing wildtype SLC38A3 indeed outperformed CAR T cells expressing this SLC38A3^mut^ variant (*p*-value: 0.016) (**Fig. S2c,d**).

As an additional validation and stringent test of CAR T cell performance, we employed a tumor rechallenge model in which mice that had cleared the primary tumor were subsequently re-injected with tumor cells in the opposite flank, without receiving any new injection of CAR T cells (**Fig. 1f**). Neither the control nor the SLC38A3^mut^ CAR T cells were able to control this second tumor, indicating insufficient CAR T cell persistence; but wildtype SLC38A3 expressing CAR T cells achieved 40% survival in this challenging model (*p*-value: 0.0012 versus control and 0.0002 versus SLC38A3^mut^) (**Fig. 1g** and **S2e**), indicating enhanced long-term persistence and efficacy of these CAR T cells.

In summary, in vivo experiments not only confirmed the enhanced antitumor efficacy conferred by overexpressing SLC38A3, but also demonstrated that this effect depends on the transporter function of SLC38A3.

### SLC38A3 improves CAR T cell stemness and metabolic fitness

We further investigated the consequences of ectopic SLC38A3 expression using immunophenotyping (**Fig. 2a**) and found increased presence of SLC38A3-expressing CAR T cells in peripheral blood (*p*-value: 0.0214) and inside the tumors (*p*-value: 0.0002) (**Fig. 2b and S3a**), with reduced expression of exhaustion markers among the tumor-infiltrating CAR T cells (**Fig. S3b,c**). While the CD4^+^/CD8^+^ ratio remained largely unchanged (**Fig. S3d**), we observed a higher frequency of memory cells among the CD4^+^ and CD8^+^ CAR T cells (*p*-value: 0.0188 and 0.0244) (**Fig. 2c**), which included CD62L^+^CD127^+^ and CD62L^+^CD45RA^-^ central memory-like T cells (**Fig. S3e**). This is likely beneficial as clinical data suggest that stem-like CAR T cells foster therapeutic success^39–41^.

**Figure 2.**
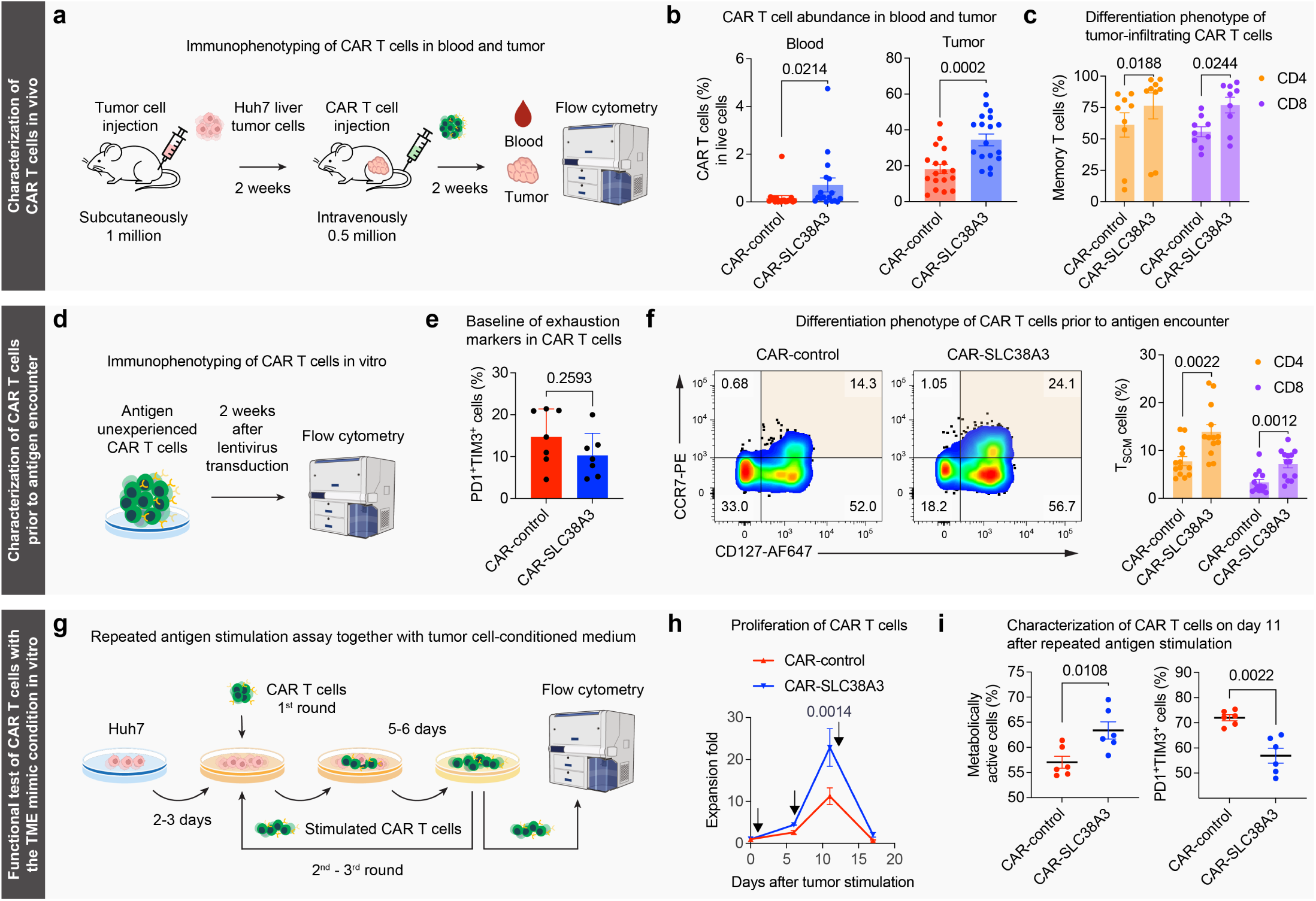
SLC38A3 enhances CAR T cell function by promoting stemness and metabolic fitness. (a) Outline of immunophenotyping for SLC38A3-expressing CAR T cells following in vivo tumor encounter. (b) Percentage of CAR T cells in blood and tumor. Each dot represents one mouse (n = 18 across five T cell donors). (c) Percentage of memory cells (CD45RA^−^) among CD4^+^ and CD8^+^ CAR T cells obtained from tumors. Each dot represents one mouse (n = 9 across three T cell donors). (d) Outline of immunophenotyping for SLC38A3-expressing CAR T cells prior to tumor encounter (in vitro). (e) Percentage of PD-1^+^TIM-3^+^ cells among CAR T cells prior to tumor encounter (n = 7 across three T cell donors). (f) Differentiation state of CAR T cells prior to tumor encounter (in vitro). Left: representative flow cytometry plots for CCR7 and CD127 expression on the CAR T cells. Right: Percentage of CCR7^+^CD127^+^ stem cell-like memory (SCM) cells among CD4^+^ and CD8^+^ CAR T cells (other markers shown in **Fig. S3f**) (n = 13 across five T cell donors). (g) Outline of the repeated antigen stimulation assay for in vitro testing of SLC38A3-expressing CAR T cells. (h) Proliferation of CAR T cells during repeated antigen stimulation with tumor cell-conditioned medium. Black arrows indicate the starting time of each stimulation round on days 0, 6, and 11 (n = 6 across three T cell donors). (i) Percentage of metabolically active cells (mitochondrial mass and cellular oxidation shown in **Fig. S3g**) and of PD-1^+^TIM-3^+^ cells among CAR T cells on day 11 of repeated antigen stimulation (n = 6 across three T cell donors). *P*-values in this figure were determined by two-tailed Mann-Whitney tests except for (h), where a two-way ANOVA with multiple-testing correction was used. Error bars show mean ± SEM.

Prior to injecting the CAR T cells into tumor-bearing mice (**Fig. 2d**), we saw no significant change in the basal expression of T cell exhaustion markers (**Fig. 2e**), indicating that the observed in vivo difference is a consequence of tumor encounter. In contrast, we detected increased expression of stem cell-like memory (SCM) markers on SLC38A3-expressing CAR T cells already prior to injection, which included elevated levels of CCR7 and CD127 (p-value: 0.0022 for CD4^+^ and 0.0012 for CD8^+^ T cells) (**Fig. 2f and S3f**), indicating that an increased stem-like memory phenotype is a constitutive, cell-intrinsic property of SLC38A3-expressing CAR T cells.

To mimic chronic antigen exposure and metabolic restrictions characteristic of solid tumors, we exposed the CAR T cells to a repeated antigen stimulation assay with supplementation of tumor cell-conditioned medium (**Fig. 2g**). By day 11, when CAR T cells typically become dysfunctional in this assay, SLC38A3-expressing CAR T cells had expanded twice as much as the control CAR T cells (*p*-value: 0.0014) (**Fig. 2h**) and maintained higher metabolic activity throughout this time-course experiment (**Fig. 2i and S3g**). We observed a higher percentage of metabolically active CAR T cells as indicated by flow cytometry with MitoTracker and CellROX (**Fig. S3g**), reduced expression of the exhaustion markers PD-1 and TIM-3 (*p*-value: 0.0022) (**Fig. 2i and S3h**) and a higher percentage of CAR T cells with a CD62L^+^CD45RO^+^ central memory phenotype (*p*-value: 0.0087) (**Fig. S3i**).

In summary, these results show that ectopic SLC38A3 expression constitutively programs CAR T cells toward a stem-like memory phenotype and enables them to sustain persistent antitumor function while mitigating T cell exhaustion, both inside the tumor (in vivo) and upon chronic antigen exposure with metabolic restriction (in vitro).

### SLC38A3 enhances amino acid uptake and glutamine metabolism in CAR T cells

We hypothesized that the enhanced metabolic fitness and antitumor efficacy of SLC38A3-expressing CAR T cells are direct consequences of the cells’ increased access to amino acids, reflecting the established role of SLC38A3 as an amino acid transporter^42^. We thus compared intracellular amino acid levels between SLC38A3-expressing and control CAR T cells (**Fig. 3a and Table S4**). We observed significantly higher cellular levels of amino acids related to glutamine metabolism upon ectopic SLC38A3 expression for both pre-infusion and tumor-infiltrating CAR T cells (**Fig. 3b and S4a-c**), indicating that SLC38A3 constitutively alters these cells’ glutamine metabolism.

**Figure 3.**
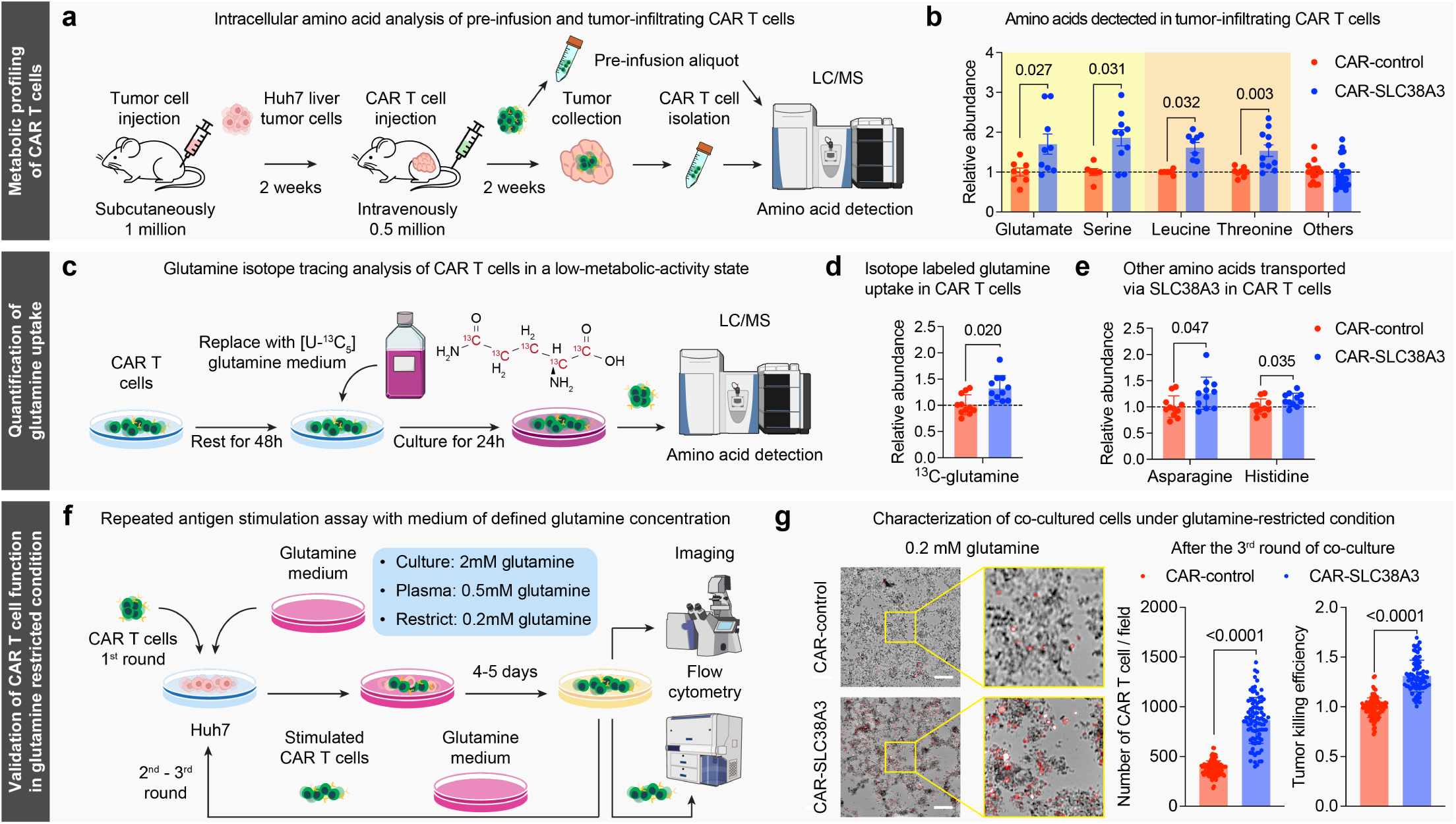
SLC38A3 improves glutamine uptake and enhances CAR T cell function under glutamine restriction. (a) Outline of intracellular amino acid profiling for pre-infusion and tumor-infiltrating CAR T cells. (b) Bar plots of intracellular amino acid levels in tumor-infiltrating CAR T cells. The yellow area (left) highlights amino acids involved in glutamine metabolism. The orange area (center) highlights essential amino acids. Aggregated results for the other amino acids are on the right (details shown in **Fig. S4c**). Each dot represents one mouse (n = 11 across four T cell donors with two to three mice each). Missing dots are due to undetected signal. (c) Outline of glutamine isotope tracing in resting CAR T cells following IL-2 removal from the culture medium. (d) Bar plot of isotope-labeled glutamine uptake in resting CAR T cells (n = 12 across four T cell donors). (e) Bar plots of asparagine and histidine levels (these two amino acids are additional substrates of SLC38A3) in resting CAR T cells (n = 12 across four T cell donors). (f) Outline of the functional testing of SLC38A3-expressing CAR T cells under defined glutamine conditions. (g) CAR T cell function after three rounds of co-culture with tumor cells under glutamine restriction. Left, representative images showing co-cultured CAR T cells (red) and tumor cells in glutamine-restricted medium (0.2 mM). Scale bars indicate 1 mm. Right, bar plots of CAR T cell numbers (fluorescent objects per image) and tumor killing efficiency (cell-free area after the detachment of dead tumor cells) (n = 60 images across three T cell donors). *P*-values in this figure were determined by two-tailed Mann-Whitney tests. Error bars show mean ± SEM.

In addition, we observed increased levels of the neutral essential amino acids leucine and threonine (**Fig. 3b**), which can be linked to glutamine metabolism: previous research discovered a role of glutamine in leucine uptake through a mechanism involving glutamine exchange for extracellular neutral amino acids, which fosters mTORC1 signaling and cell growth^43,44^. Our data thus suggest a putative mechanism by which SLC38A3 enhances broader metabolic activity and amino acid uptake in support of CAR T cell function inside solid tumors.

To validate glutamine uptake as the cause of these metabolic changes, we performed stable-isotope tracing using [U-^13^C_5_]-glutamine in resting CAR T cells (**Fig. 3c and Table S5**), which was selected for their low metabolic activity and minimal confounding by de novo amino acid synthesis. We found that the higher abundance of intracellular glutamine in SLC38A3-expressing CAR T cells compared to controls (*p*-value: 0.02) was predominantly driven by the ^13^C-labeled isotopologue acquired from the extracellular medium (**Fig. 3d and S4d**), reinforcing that ectopic SLC38A3 expression directly increases glutamine uptake in CAR T cells.

As expected for resting CAR T cells, we observed minimal incorporation of glutamine-derived ^13^C into downstream catabolic intermediates and little difference in the overall amino acid profiles (**Fig. S4d,e**). Nevertheless, asparagine and histidine levels were increased in SLC38A3-expressing CAR T cells (*p*-value: 0.047 and 0.035) (**Fig. 3e**). Considering the slow metabolic turnover of resting cells and the status of histidine as an essential amino acid, this is likely a direct consequence of SLC38A3’s ability to import not only glutamine, but also asparagine and histidine^42^. Our results thus indicate SLC38A3-mediated transport of its three substrates in CAR T cells.

In summary, these experimental data support a functional role of ectopic SLC38A3 expression in enhanced cellular uptake of several amino acids that have well-established roles in supporting T cell proliferation and activity.

### SLC38A3 sustains CAR T cell function under glutamine restriction

Glutamine restriction is common among solid tumors and a challenge for infiltrating T cells^45–47^. To test whether ectopic SLC38A3 expression can help maintain CAR T cell function under glutamine restriction, we performed repeated antigen stimulation experiments using media with defined glutamine concentrations: standard cell culture level (2 mM), physiological plasma level (0.5 mM), and nutrient-restricted TME-like level (0.2 mM) (**Fig. 3f**).

After the first exposure to tumor cells, SLC38A3-expressing and control CAR T cells achieved comparable proliferation and antitumor efficacy across all three glutamine levels (**Fig. S4f**). However, after the second exposure to tumor cells, the SLC38A3-expressing CAR T cells showed significantly higher proliferation for glutamine’s physiological plasma level (*p*-value: 0.0012) (**Fig. S4g**); and we observed significantly higher expression of the stemness marker CD62L when aggregating across all three glutamine levels (*p*-value: 0.012) (**Fig. S4h**).

After the third exposure to tumor cells, control CAR T cells cultured in medium with the lowest, TME-like glutamine level had much lower survival (*p*-value < 0.0001) and antitumor efficacy (*p*-value < 0.0001) compared to SLC38A3-expressing CAR T cells (**Fig. 3g**). This difference was less pronounced for higher glutamine levels, where the control CAR T cells remained partially effective (**Fig. S4i**). These results further support that enhanced glutamine uptake conferred by ectopic expression of SLC38A3 fosters CAR T cell function under glutamine restriction.

In summary, our observations indicate that the increased access to extracellular glutamine conferred by ectopic SLC38A3 expression is particularly relevant for sustained CAR T cell function upon extended antigen exposure with simultaneous glutamine restriction – which is a typical challenge that CAR T cells face in solid tumors.

### SLC38A3 enhances CAR T cell efficacy across tumor models with high glutamine consumption

To validate and generalize the beneficial effect of ectopic SLC38A3 expression in CAR T cells targeting glutamine-restricted tumors, we complemented the Huh7 liver tumor model used in our previous experiments with a second widely used liver tumor model (HepG2) and an ovarian tumor model (SKOV3). Metabolomic profiling of tumor cell-conditioned medium by mass spectrometry (**Fig. 4a and Table S6**) confirmed that all three are strong consumers of glutamine, while we observed differences in terms of other metabolic characteristics (**Fig. 4b and S5a,b**).

**Figure 4.**
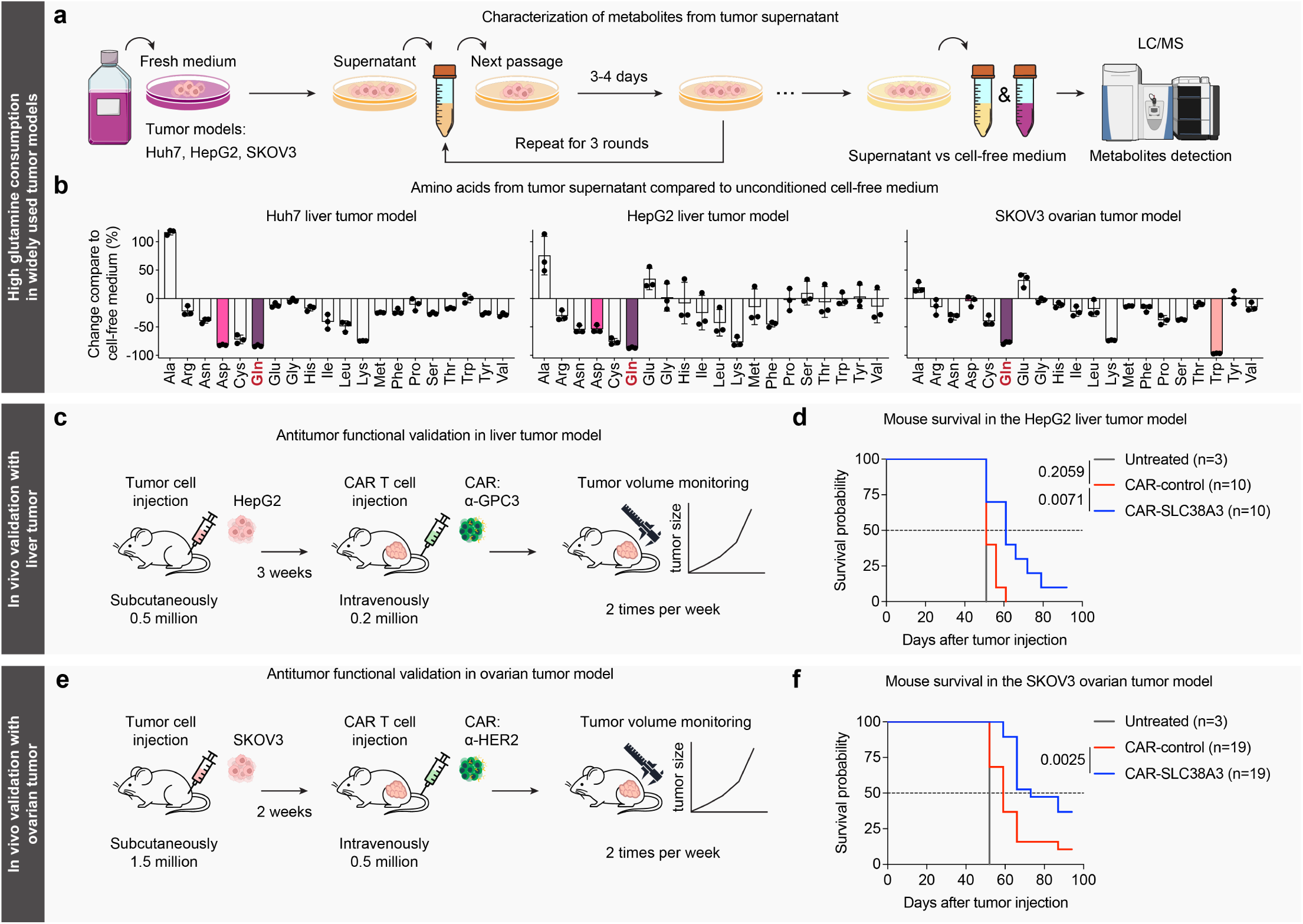
SLC38A3 enhances CAR T cell efficacy in several tumor models with high glutamine consumption. (a) Outline of amino acid profiling applied to tumor cell-conditioned medium (supernatant) of Huh7, HepG2, and SKOV3 tumor cells. (b) Bar plots of amino acid changes in tumor cell-conditioned medium compared to unconditioned cell-free medium (n = 3). (c) Outline of in vivo validations for SLC38A3-expressing CAR T cells in xenograft liver tumors (HepG2). (d) Survival curves for HepG2 tumor-bearing mice left untreated (grey), injected with control CAR T cells (red), or with SLC38A3-expressing CAR T cells (blue). Experiments were performed independently for two T cell donors with five mice each. (e) Outline of in vivo validations for SLC38A3-expressing CAR T cells in xenograft ovarian tumors (SKOV3). (f) Survival curves for SKOV3 tumor-bearing mice left untreated (grey), injected with control CAR T cells (red), or with SLC38A3-expressing CAR T cells (blue). Experiments were performed independently across four T cell donors with five mice each. *P*-values in (d) and (f) were determined by the log-rank (Mantel-Cox) test. Error bars in (b) show mean ± SD.

When we treated HepG2 tumor-bearing mice with a deliberately non-curative dose of CAR T cells targeting GPC3 (**Fig. 4c**), the control CAR T cells barely extended survival compared to untreated mice (*p*-value: 0.2059); in contrast, the same number of SLC38A3-expressing CAR T cells provided a substantial survival advantage compared to the control CAR T cells (*p*-value: 0.0071) (**Fig. 4d and S5c**).

We obtained similar results for the ovarian tumor model and CAR T cells targeting a different tumor antigen (HER2) (**Fig. 4e**). SKOV3 tumor-bearing mice treated with SLC38A3-expressing CAR T cells were more than twice as likely to survive to day 70 than mice treated with control CAR T cells (*p*-value: 0.0025) (**Fig. 4f and S5d**).

Finally, we assessed the durability of the antitumor response in the SKOV3 tumor model with a tumor rechallenge experiment similar to the one presented in **Figure 1f**. Remarkably, most mice previously treated with SLC38A3-expressing CAR T cells were able to reject the tumor rechallenge (3 million SKOV3 cells injected into the opposite flank) and survived, while the same rechallenge was universally lethal for the mice treated with control CAR T cells (*p*-value: 0.0185) (**Fig. S5e,f**), indicating longer-lasting protection by SLC38A3-expressing CAR T cells.

In summary, we found that SLC38A3 expression enhances the therapeutic efficacy of CAR T cells in three widely used solid tumor xenograft models that are characterized by high glutamine consumption, thus validating the concept of enhancing the metabolic fitness of CAR T cells through ectopic nutrient transporter expression.

## Discussion

To stay active inside solid tumors, CAR T cells compete with the cancer cells for scarce nutrients^48,49^. Adding to the challenge, cancer cells frequently evolve mechanisms that optimize their ability to take up nutrients^50–52^, which aggravates nutrient depletion and contributes to the highly immunosuppressive TME that CAR T cells encounter in many solid tumors. Here, we pursued in vivo gain-of-function screening as a form of “artificial evolution” that seeks to enhance the CAR T cells’ ability to compete with tumor cells for access to critical nutrients.

We developed the CARGO method for overexpression screens in xenograft tumor models and human primary CAR T cells, and tested the in vivo effects of overexpressing 452 solute carrier (SLC) transmembrane transporters in a liver tumor model that reflects the hostile TME of human hepatocellular carcinoma. Our method incorporates efficient barcode sequencing and UMI-assisted quality control; it addresses key limitations of CRISPR activation screening, which struggles to achieve robust overexpression across all target genes^27,29^; and it was successfully validated by deep-coverage in vivo screening for one of the largest membrane protein superfamilies in the genome.

From our screening hits, we selected SLC38A3 as the most promising “metabolic booster” of CAR T cell function. SLC38A3 is not endogenously expressed in T cells but highly expressed in brain and liver^53,54^, making it a prime example of our hypothesis that gain-of-function screening can effectively repurpose genes from other biological contexts to enhance CAR T cells. We performed extensive in vivo validations, observing superior antitumor efficacy and prolonged survival with SLC38A3-expressing CAR T cells across three widely used xenograft tumor models (Huh7, HepG2, SKOV3), two tumor types (hepatocellular and ovarian carcinoma), and two target antigens (GPC3, HER2). We also showed that the beneficial effect of ectopic SLC38A3 expression depends on its transporter function (the transport-deficient SLC38A3^mut^ variant was ineffective), and we showed that SLC38A3 not only increases short-term efficacy but also long-term persistence and function in tumor rechallenge experiments.

We found that ectopic SLC38A3 expression induces a stem-like memory phenotype in CAR T cells already prior to antigen encounter. It also helps CAR T cells maintain metabolic fitness and reduces exhaustion upon chronic antigen exposure and under metabolic restriction. Consistent with SLC38A3’s function as a glutamine transporter, we showed by [U-^13^C_5_]-glutamine tracing that SLC38A3 enhances the CAR T cells’ uptake of glutamine, a critical fuel for oxidative phosphorylation^55,56^ and a metabolic requirement for T cell growth and proliferation^56^.

Notably, recent evidence suggests that T cells rely more heavily on glutamine than glucose to replenish the tricarboxylic acid (TCA) cycle and to provide biosynthetic substrates in vivo^57^. At the same time, many aggressive tumors exhibit a high dependency on glutamine^47,58^ and upregulate transporters such as SLC1A5 and SLC38A1/2 to bolster their access to glutamine^59,60^. In this competition for glutamine, ectopic expression of SLC38A3 confers a competitive advantage to the CAR T cells, explaining the superior in vivo antitumor performance that we observed.

Beyond the presented in vivo screening method and our discovery of SLC38A3 as a booster of CAR T cell function in glutamine-restricted tumors, we also noted a striking discrepancy between our initial in vitro screens (**Supplementary Note**) and subsequent in vivo screens. For example, in vitro screening identified mitochondrial pyruvate carrier 1 and 2 (MPC1 and MPC2) as top hits for enhancing CAR T cell expansion under repeated tumor stimulation. This is a biologically plausible finding as these two genes may help CAR T cells leverage lactate oxidation for sustained antitumor activity^61^. However, their effect was not recapitulated in our in vivo screening and did not validate in dedicated in vivo experiments. Our study thus highlights the limitations of standard in vitro assays for modeling the metabolic challenges in the TME and underlines the importance of screening in suitable in vivo models.

Our study has certain limitations that warrant consideration. First, while we used cell line-derived xenograft models to facilitate large in vivo screens, our method can also support screens in patient-derived xenografts. Second, our focus on xenograft models of human tumors enabled in vivo screens with human primary CAR T cells, but working with immunodeficient mice removes host immune cells from the TME. Third, glutamine restriction is a common but not a universal property of solid tumors^62,63^, and the overexpression of other SLCs may be more beneficial for tumors with different metabolic properties – pointing to the need for a “precision immunometabolism” approach that tailors CAR T cells to the metabolic properties of the tumors that they are intended to treat.

In conclusion, this study establishes ectopic expression of SLC transmembrane transporters as an effective way of enhancing CAR T cell function in the nutrient-deprived TME of solid tumors, and it provides an in vivo screening method enabling the systematic discovery and optimization of metabolic boosters for cell-based immunotherapy.

## Methods

### Experimental models

#### Mice

NSG mice (NOD.Cg-*Prkdc*^scid^*Il2rg*^tm1Wjl^/SzJ) were bred and maintained under specific-pathogen-free conditions at the Core Facility Laboratory for Animal Breeding and Husbandry of the Medical University of Vienna, which is authorized for animal breeding and experimentation under license BMWFW-66.009/0403-WF/V/3b/2014. Experiments followed established guidelines and were approved by the institutional ethical committee at the Department for Biomedical Research of the Medical University of Vienna according to license BMWFW-2020-0.605.586 granted by the Austrian Federal Ministry of Education, Science and Research (BMBWF). The animal experiments were performed in a blinded manner to reduce potential biases. The determination of group sizes and the power calculation were carried out according to the approved animal research license.

#### Cancer cell lines

The hepatocellular carcinoma cell lines Huh7 (JCRB #JCRB0403) and HepG2 (ATCC #HB-8065) as well as the ovarian carcinoma cell line SKOV3 (ATCC #HTB-77) were used as tumor models. Lenti-X 293T cells (TaKaRa #632180) were used for lentivirus production. Cell lines were cultured in high-glucose DMEM (Sigma #D5796) supplemented with 10% FBS (Sigma #F7524) and 1% penicillin–streptomycin (Gibco #15140122). Cells were split and culture medium was exchanged every 3-4 days following standard procedures for adherent cells. Cell lines were validated by short tandem repeat loci profiling (Microsynth). All cell lines used in this study were negative for Mycoplasma contamination throughout the study; this was confirmed by regular PCR-based testing.

#### Primary T cells

Peripheral blood samples from healthy donors were obtained through the Austrian Red Cross as blood packs with buffered sodium citrate as anticoagulant. Informed consent was obtained from all sample donors. The research complied with all relevant ethical regulations for working with human samples and was approved by the ethical committees of the contributing institutions (Austrian Red Cross and the Medical University of Vienna). Human primary T cells were isolated using RosetteSep Human T Cell Enrichment Cocktail (Stemcell #15061) to purify CD3**^+^** T cells. After incubation with the antibody cocktail, cells were centrifuged using Lympho-prep (Stemcell #7861) in SepMate-50 tubes (Stemcell #85460), washed and treated with 1× RBC Lysis Buffer (eBioscience #00-4333-57). Isolated T cells were immediately stimulated and cultured, or frozen in liquid nitrogen in single-use aliquots. T cell culture used CTS OpTmizer T-Cell Expansion Serum Free Medium (Gibco #A1048501), 1000 ml of which was supplemented with 26 ml of CTS OpTmizer T-Cell Expansion Supplement, 1% L-Glutamine (200 mM, Gibco #25030081), and 1% penicillin–streptomycin (Gibco #15140122). The medium was supplemented with 100 U/ml IL-2 (Stemcell #78145.2) to support T cell growth. T cell density was monitored by cell counting and adjusted to 0.5 million cells/ml at each passage. Medium and IL-2 were replenished every 2-3 days.

### Screening vector and library

#### Cloning of the CARGO vector for in vivo screening

We devised the CARGO vector (“CAR + Gene Overexpression”) as a third-generation lentiviral vector that enables simultaneous delivery of a CAR and a payload gene or an ORF library into human primary T cells. The vector backbone was based on the pRRLSIN.cPPT.PGK-GFP.WPRE plasmid (Addgene #12252). To coordinate the co-expression of reporter gene (GFP) and the payload gene or ORF library while maintaining a small vector size, we replaced the hPGK promoter with a bidirectional promoter cassette similar to a previous study^64^.

We linearized the vector backbone by digestion with SalI (NEB #R3138S) and XhoI (NEB #R0146S) restriction enzymes and assembled it with a PCR fragment containing the bidirectional gene expression cassette via Gibson assembly using NEBuilder HiFi DNA Assembly Master Mix (NEB #E2621S). Specifically, this cassette consists of a hEF1α promoter for high gene expression followed by a multiple cloning site for gene insertion, and a minimal CMV promoter in the opposite orientation driving GFP expression used as a reporter for transduction.

We then cloned the selected CAR into the CARGO vector. Two CAR constructs were used in this study, targeting GPC3 (hepatocellular carcinoma) and HER2 (ovarian carcinoma). Their scFv fragments are derived from GC33^65^ and 4D5^66^ antibodies for specific targeting of the GPC3 and HER2 antigens. Both CAR receptors contain an Nterminal CD8a signal peptide (MALPVTALLLPLALLLHAARP) for membrane targeting, and a Myc-tag (EǪKLISEEDL) for CAR staining. The CD8a hinge-transmembrane region (Uniprot: P01732-1, aa 138-206) was used to connect the cytoplasmic region of costimulatory domain CD28 (Uniprot: P10747-1, aa 180-220) followed by CD3ζ intracellular domain (Uniprot: P20963-1, aa 52-164). Both CAR sequences were codon optimized, synthesized as gBlocks (IDT), and cloned into the CARGO vector at the multiple cloning site (BamHI and SalI) via Gibson assembly.

To create a landing site for the payload gene, we introduced a T2A self-cleaving peptide downstream of the CAR sequence followed by two AarI restriction sites as the multiple cloning site. To enable effective barcode sequencing for the ORF library, an 80 bp filler sequence followed by an Illumina i7 sequence was introduced downstream of the multiple cloning site. The truncated nerve growth factor receptor (tNGFR) was used as control vector payload as described previously^23,67–69^.

The ORF library comprising codon-optimized cDNAs for 452 SLCs (Addgene #213694) was amplified using a common forward primer and individual reverse primers, each containing a unique 10 bp barcode and 6 random nucleotides used as UMIs (**Table S7**). PCR products were pooled, purified using 1× vol AmpliClean magnetic beads (Nimagen #AP-050), and cloned into the CARGO vector using the GeneBuilder cloning kit (GenScript #L00701-50).

#### Lentivirus production and transduction

Lenti-X cells were seeded in 15 cm dishes with lentivirus production medium consisting of Opti-MEM (Gibco #51985026), 5% FBS, and 200 µM sodium pyruvate (Gibco #11360-070). After cell growth overnight, cells were transfected with 27 µg of the transfer plasmid, 13 µg of the packaging plasmid pMDLg/pRRE (Addgene #12251), 13 µg of the pRSV-Rev (Addgene #12253), and 13 µg of the pMD2.G (Addgene #12259), using Lipofectamine 3000 (Invitrogen #L3000150). Exchange to fresh lentivirus production medium was performed after 6 hours. Supernatant was harvested at 24 and 48 hours, respectively. The supernatant was filtered through a 0.45 µm filter, and the virus was concentrated 100× using Lenti-X concentrator (Takara #631232) following the manufacturer’s protocol. The concentrated virus was resuspended in PBS and frozen at -80 °C in single-use aliquots.

For human T cell transduction with lentivirus, T cells were first stimulated with ImmunoCult Human CD3/CD28 T Cell Activator (Stemcell #10991) following the manufacturer’s protocol. 24 hours after stimulation, concentrated lentivirus was titrated and applied to cells with transduction efficiency of 20-40% across three donors to minimize the chance of multiple SLCs being introduced into the same cell. After another 16 hours, the virus medium was completely replaced with a 4-fold larger volume of the complete T cell culture medium.

#### Library amplification and barcode sequencing

Genomic DNA from pre-infusion or tumor-infiltrating CAR T cells was purified using the ǪIAamp DNA Mini Kit (Ǫiagen #51306), and ORF library barcodes were amplified using custom primers in two consecutive PCR reactions. In the first PCR, staggered forward primers introduced the Illumina i5 sequence, while the reverse primer was designed to bind downstream of the Illumina i7 sequence in the vector backbone, resulting in PCR products containing both the i5 and i7 sequences. In the second PCR, with the first PCR product as template, Illumina adapters and multiplexing indices were introduced. Independently, plasmid library sequencing was performed by using 1 ng of library plasmid as PCR template. All PCRs were performed using Ǫ5 high fidelity polymerase (NEB #M0491L). Staggered ORF library barcode amplification primers (first PCR) and Illumina barcoding primers for the ORF library (second PCR) are provided in **Table S7**. Barcode amplicons generated by the second PCR were multiplexed and sequenced on an Illumina NovaSeq 6000 instrument by the Biomedical Sequencing Facility at CeMM.

### Xenograft tumor models

#### In vivo screening

The in vivo screening was performed using the xenograft Huh7 liver tumor mouse model. NSG mice of both sexes at 8-12 weeks of age were subcutaneously injected with 3 million Huh7 cells suspended in 150 µl PBS into the right flank. After 2 weeks (corresponding to tumor volumes of approximately 300-500 mm^3^), mice were treated with 2 million CAR-SLC library T cells in 100 µl PBS by intravenous injection. After another 2 weeks, mice were sacrificed with CO_2_, followed by collection and dissociation of the tumors. The CAR T cells were isolated from the tumor cell suspension using negative selection with the EasySep Dead Cell Removal Kit (Stemcell #17899) followed by positive selection using the EasySep Release Human CD3 Positive Selection Kit (Stemcell #17751). Cell pellets were snap-frozen in a liquid nitrogen container and stored at -80 °C until DNA extraction.

#### Antitumor functional test of CAR T cells in vivo

NSG mice of both sexes at age of 8-12 weeks were subcutaneously injected with the tumor cells (1 million Huh7, 0.5 million HepG2, or 1.5 million SKOV3) suspended in 100 µl PBS into the right flank. After 2 weeks (Huh7, SKOV3) or 3 weeks (HepG2), corresponding to tumor volumes of approximately 150-300 mm^3^ for Huh7 and HepG2, or 50 mm^3^ for SKOV3, mice were treated with the CAR T cells (0.5 million for the Huh7 and SKOV3 models and 0.2 million for the HepG2 model) in 100 µl PBS by intravenous injection. Tumor size was monitored twice per week throughout the experiment using a caliper to measure tumor length (*L*; greatest longitudinal measurement) and width (*W*; greatest transverse measurement). Tumor volumes (*V*) were calculated using the formula: *V* = (*L* × *W*^2^) / 2. Mice were sacrificed when the tumors reached the maximum size specified in the animal research license.

For characterization of CAR T cells in vivo, tumor-bearing mice were sacrificed 2 weeks after CAR T cell treatment. Tumor and peripheral blood samples were collected, and tumors were dissociated to isolate CAR T cells for the indicated profiling. For the tumor rechallenge model (initial dose of 1 million Huh7 or 1.5 million SKOV3), when tumor-bearing mice were assessed as tumor-free for at least one month after CAR T cell treatment, 2 million Huh7 cells or 3 million SKOV3 cells were injected into the left flank to inoculate the rechallenge tumor.

#### Tissue dissociation

Blood sampling was performed by tail-vein puncture or cardiac puncture of the mice. Blood was collected in the Microtainer tube containing K_2_ EDTA (BD #10346134) and kept at room temperature until further processing. Tumor sampling was performed with gentleMACS Octo Dissociator (Miltenyi Biotec #130-096-427) and Tumor Dissociation Kit (Miltenyi Biotec #130-095-929) according to the manufacturer’s instructions. For both blood and tumor tissue, the cells were treated with RBC Lysis Buffer (BioLegend #420302) to remove red blood cells and filtered through a 40-μm cell strainer (Falcon #352340) before further processing by flow cytometry or FACS.

### In vitro experiments

#### Flow cytometry and cell sorting

For cell surface staining, cells were incubated with antibodies at 4 °C for 30 min in Cell Staining Buffer (BioLegend #420201). DAPI (Thermo Scientific #62248) was used to exclude dead cells. The expression of SLC38A3 was intracellularly detected by staining the HA-tag fused to the N-terminus. CAR T cells were fixed and permeabilized using Cyto-Fast Fix/Perm Buffer Set (BioLegend #426803) following the manufacturer’s protocol for intracellular staining. To stain mitochondria and activation-induced ROS, CAR T cells were incubated with 20 nM MitoTracker Orange (Invitrogen #M7510) and 5 μM CellROX Deep Red (Invitrogen #C10422) in a cell incubator for 20 min. Ǫuantification of CAR T cell numbers was performed by flow cytometry-based bead counting using Precision Count Beads (BioLegend #424902) following the manufacturer’s protocol. All flow cytometry profiling was performed using a BD LSRFortessa Cell Analyzer. Data were analyzed with FlowJo v10.8.1 (Tree Star). Cell sorting was performed on a Sony SH800S cell sorter. All flow cytometry and cell sorting antibodies are listed in **Table S8**.

#### Repeated antigen stimulation assay

Repeated antigen stimulation was used to model chronic tumor stimulation in vitro. Huh7 cells were first cultured alone until confluence. CAR T cells were then seeded and co-cultured with tumor cells in a 1:3 effector cell to target cell (E:T) ratio. For the co-culture, OpTmizer T cell medium conditioned by Huh7 cells was used in **Fig. 2g** to mimic the metabolically restricted TME; and OpTmizer T cell medium supplemented with defined glutamine concentrations was used in **Fig. 3f** to test CAR T cell function under glutamine-restricted conditions. After coculture for the indicated duration, CAR T cells were counted and seeded again for repeated stimulation until too few CAR T cells could be collected for the next round of stimulation.

The repeated antigen stimulation assay was also used for the in vitro screening (**Supplementary Note**). There, a total of 3 million CAR-SLC library T cells were co-cultured with confluent Huh7 cells in T-175 flasks with complete CTS OpTmizer T cell media, freshly added prior to each round of stimulation. CAR T cells were counted and reseeded weekly with an aliquot cryopreserved in the meantime for subsequent barcode sequencing. This process was repeated for 8 weeks until fewer than 3 million CAR T cells could be harvested for the next round of co-culture.

#### Live cell imaging

Live cell imaging was performed to visualize the co-culture of CAR T cells and tumor cells. CAR T cells and Huh7 tumor cells were co-cultured as shown in **Fig. 3f** using media with defined glutamine concentrations. By the end of the experiment, cells were imaged using the Revvity Opera Phenix high-content confocal imaging system equipped with a 20× objective. For each condition, 6-10 fields of view were imaged per well of a 6-well plate format. Primary image analysis and quality control were performed using the Harmony software v5.3. Subsequent quantitative image analysis was performed using Fiji v2.9.0. Images were first scaled to ensure consistent spatial calibration across acquisitions. Each image was subsequently converted to a binary format, and intensity thresholds were determined empirically for each channel to ensure accurate segmentation. Surface area measurements were obtained using the “Set Measurements” function, with area selected as the parameter of interest, followed by measurement of the threshold regions. CAR T cell puncta quantification was performed using the “Analyze Particles” function, with size and circularity gates applied to exclude artifacts and debris. All quantification parameters were kept constant across conditions and donors within an experiment.

### Profiling assays

#### Metabolite analysis of tumor supernatant

Huh7, HepG2, and SKOV3 tumor cells were cultured in the complete DMEM medium and passaged every 3-4 days upon reaching confluence. Supernatant was collected and reused in the next passage. After three rounds of passaging, tumor supernatant and unconditioned cell-free medium in the incubator were collected for hydrophilic metabolite analysis. 20 µl of medium was mixed with internal standards and 300 µl methanol, incubated for 20 min at 450 rpm at room temperature, and centrifuged for 10 min at 10,000 × g at 4 °C. The supernatant was transferred to a vial with insert and gently evaporated. The samples were then reconstituted in 100 µl HPLC-grade water and subjected to LC-ESI-MS/MS analysis. Targeted metabolomics profiling (METAB02 method) was performed by the Metabolomics Facility at CeMM. The PeakBot software (v0.9.54) was used for data processing^70^. Ten-point calibration curves with internal standardization were constructed for absolute quantification of metabolites.

#### ^13^C-metabolite labeling and intracellular amino acid analysis

CAR T cells were rested by culturing in CTS OpTmizer T-Cell medium without IL-2 for 48 h. They were then washed with PBS and resuspended in CTS OpTmizer T-Cell medium (no glutamine is included in this medium) supplemented with 0.68 mM [U-¹³C_5_]-L-Glutamine (Cambridge Isotope Laboratories #CLM-1822) and cultured another 24 h. After incubation, cell pellets were collected and transferred into 1.5 ml Eppendorf tubes, immediately snap-frozen in a liquid nitrogen container, and stored at -80 °C. Cell pellets were then spiked with 20 µl of an isotopically labeled internal standards mixture. Metabolites were extracted by adding 400 µl of ice-cold 80% methanol, followed by ultrasound-assisted extraction for 5 min with samples maintained on ice. Extracts were then centrifuged for 10 min at 10,000 × g and 4 °C. Supernatants were collected and evaporated to dryness under a gentle nitrogen stream at room temperature. Dried extracts were reconstituted in 20 µl of methanol prior to LC-MS analysis.

Liquid chromatography mass spectrometry (LC-MS) profiling was performed using a Vanquish UHPLC system coupled to an Orbitrap Ǫ Exactive mass spectrometer (Thermo Scientific). Chromatographic separation was performed using an ACǪUITY UPLC BEH Amide analytical column (1.7 µm, 2.1 × 100 mm; Waters) equipped with a VanGuard BEH Amide pre-column (2.1 × 5 mm; Waters). The column temperature was maintained at 40 °C, and the injection volume was 10 µl. The mobile phases consisted of: (A) water containing 0.15% (v/v) formic acid and 10 mM ammonium formate, and (B) 85% acetonitrile (v/v) containing 0.15% (v/v) formic acid and 10 mM ammonium formate. Gradient elution was performed at a flow rate of 0.4 ml/min with a total run time of 17 min. The Orbitrap Ǫ Exactive mass spectrometer was operated in positive electrospray ionization mode with the following parameters: spray voltage, 3.5 kV; capillary temperature, 350 °C; auxiliary gas heater temperature, 400 °C; sheath gas flow rate, 50; auxiliary gas flow rate, 12. Data were acquired in full MS mode over an m/z range of 50–400 at a resolution of 35,000, with an AGC target of 1×10⁶ and a maximum injection time of 50 ms. Metabolite quantification was performed using TraceFinder 4.1 software (Thermo Scientific). An eight-point linear calibration curve with internal standardization and 1/*x* weighting was used for quantification.

### Bioinformatic data analysis

#### Barcode sequencing analysis

For barcode sequencing analysis, 5’ flanking regions (ACCCAGCTTTCTTGTACAAAG) and 3’ downstream flanking regions (TGCATAGCTGAGCTACTTGA) were trimmed from the reads using Cutadapt and the remaining sequences were trimmed to 10 bp to obtain the ORF barcodes. Occurrences of each barcode were counted for each sample, and significant enrichment or depletion was determined by comparing barcode counts between pre-infusion and tumor-infiltrating CAR T cells using DESeq2 (Version 1.40.2) with default parameters.

#### Unique molecular identifier analysis

To quantify the unique molecular identifier (UMI) diversity for each ORF barcode (**Table S3**), the aligned sequencing reads were parsed using Pysam, and read sequences were scanned using the Python regex library to extract the 10 bp ORF barcode (**Table S1**) as well as the adjacent 6 bp UMI. This pattern-matching allowed for up to one sequence mismatch in the 5’ flanking regions (ACCCAGCTTTCTTGTACAAAG). The 10 bp barcodes were matched to their respective target genes requiring an exact match to the reference barcode library. To evaluate UMI stability for pre-infusion and tumor-infiltrating CAR T cells, the unique UMI sequences per gene were counted, averaged across all replicates of each donor, and visualized for each donor and condition using violin plots.

#### Public RNA-seq data analysis

To assess SLC gene expression in tumor cells, T/NK cells, and other cell types from liver tumors, a publicly available single-cell RNA-seq dataset^37^ (GSE149614) was analyzed using Scanpy^71^. Cells with fewer than 100 detected genes and genes expressed in fewer than 10 cells were excluded, followed by total-count normalization and log1p-transformation. Fibroblasts and endothelial cells were combined into “Stromal cells”; and B cells, myeloid cells, and TAMs were combined into “Other immune” cells. Expression of SLC genes in the indicated cell types across tumor and normal regions was visualized with bubble plots using the sc.pl.dotplot function of Scanpy.

#### Isotopologue analysis and flux calculation

Mass distribution vectors were extracted from raw ion chromatograms using MATLAB. For each metabolite, extracted ion chromatograms corresponding to all detectable isotopologues (M+0 to M+n, where n is the number of carbon atoms in the metabolite) were integrated, normalized to the total isotopologue signal, and corrected for the contribution of naturally occurring ^13^C isotopes using a matrix-based algorithm^72^. Corrected isotopologue abundances were used to calculate the fractional contribution (FC) of carbon originating from the labeled substrate, representing the average fraction of metabolite carbon derived from the tracer across the complete isotopologue distribution: FC = (Σᵢ₌₀ⁿ i × mᵢ)/(n × Σᵢ₌₀ⁿ mᵢ), where *n* is the number of carbon atoms in the metabolite, *i* denotes the isotopologue index, and *m_i_* denotes the abundance of the isotopologue *i*. Each isotopologue was weighted according to its labeling state, providing a quantitative estimate of tracer-derived carbon incorporation into the metabolite pool. FC values ranged from 0 (no incorporation of labeled carbon) to 1 (all carbon atoms within the metabolite pool are derived from the labeled substrate)^73^. Glutamine uptake was determined by quantifying the increased intracellular ^13^C-labeled glutamine within the whole glutamine pool after 24 h of incubation.

### Statistical analysis

Statistical analysis and data plotting were performed with GraphPad Prism v11. Statistical comparisons between two groups were performed using the two-tailed Mann-Whitney U test. Statistical tests comparing multiple groups with two factors were performed by two-way ANOVA followed by Šídák’s multiple comparisons test. Statistical tests comparing metabolite abundance in tumor supernatants were performed using multiple unpaired t tests followed by two-stage step-up multiple comparisons test. Statistical tests for the survival curves were performed using the log-rank (Mantel-Cox) test. *P* values below 0.05 were considered statistically significant. The specific statistical tests, numbers of replicates, and error bar definitions are indicated in each figure legend.

## Materials availability

The CARGO vector (“CAR + Gene Overexpression”) for in vivo gain-of-function screening in CAR T cells has been deposited for distribution by Addgene (#261910). The ORF library comprising 452 codon-optimized SLC transmembrane transporters is also provided through Addgene (#213694).

## Data availability

Details of the SLC ORF library including gene names, barcodes, and DNA sequences are provided in **Table S1**. The results of the in vivo screening are provided in **Table S2** and the UMI counts are provided in **Table S3**. Intracellular amino acid data for pre-infusion and tumor-infiltrating CAR T cells are provided in **Table S4**. Amino acid data for ^13^C-glutamine tracing assay in resting CAR T cells are provided in **Table S5**. Metabolomic data for tumor cell-conditioned medium are provided in **Table S6**. Primer sequences for library cloning and barcode sequencing are provided in **Table S7**. Flow cytometry antibodies used for immunophenotyping are listed in **Table S8**. In vitro screening data discussed in the **Supplementary Note** are provided in **Table N1**.

## Acknowledgements

We thank the Austrian Red Cross for providing blood samples of healthy donors; the Biomedical Sequencing Facility at CeMM for assisting with next generation sequencing; the Metabolomics Facility at CeMM for metabolomics profiling and data analysis; the Core Facility Laboratory Animal Breeding and Husbandry of the Medical University of Vienna for hosting the animal work and providing access to materials, services, and equipment; and the members of the Bock lab for advice and discussions. This study was co-funded by an EMBO fellowship (grant no. ALTF 1163-2021) and an Austrian Science Fund (FWF) ESPRIT fellowship (10.55776/ESP4624425) to WL; an FWF Special Research Area grant (SFB F70) to CB and TK; and an ERC Consolidator Grant (grant agreement no. 101001971) of the European Union’s Horizon 2020 research and innovation program to CB.

## Author contributions

Conceptualization: WL, GW, GSF, CB; Methodology: WL, GW; Investigation: WL, GW, JL, NP, RSF, MJ, CG, FP, SL; Resources: GSF; Formal analysis: WL, GW, RB, RSF; Visualization: PM; Writing – Original Draft: WL, TK, CB; Writing – Review & Editing: All authors; Supervision: LdR, GSF, CB; Funding Acquisition: WL, CB.

## Supplementary Figures

**Figure S1.**
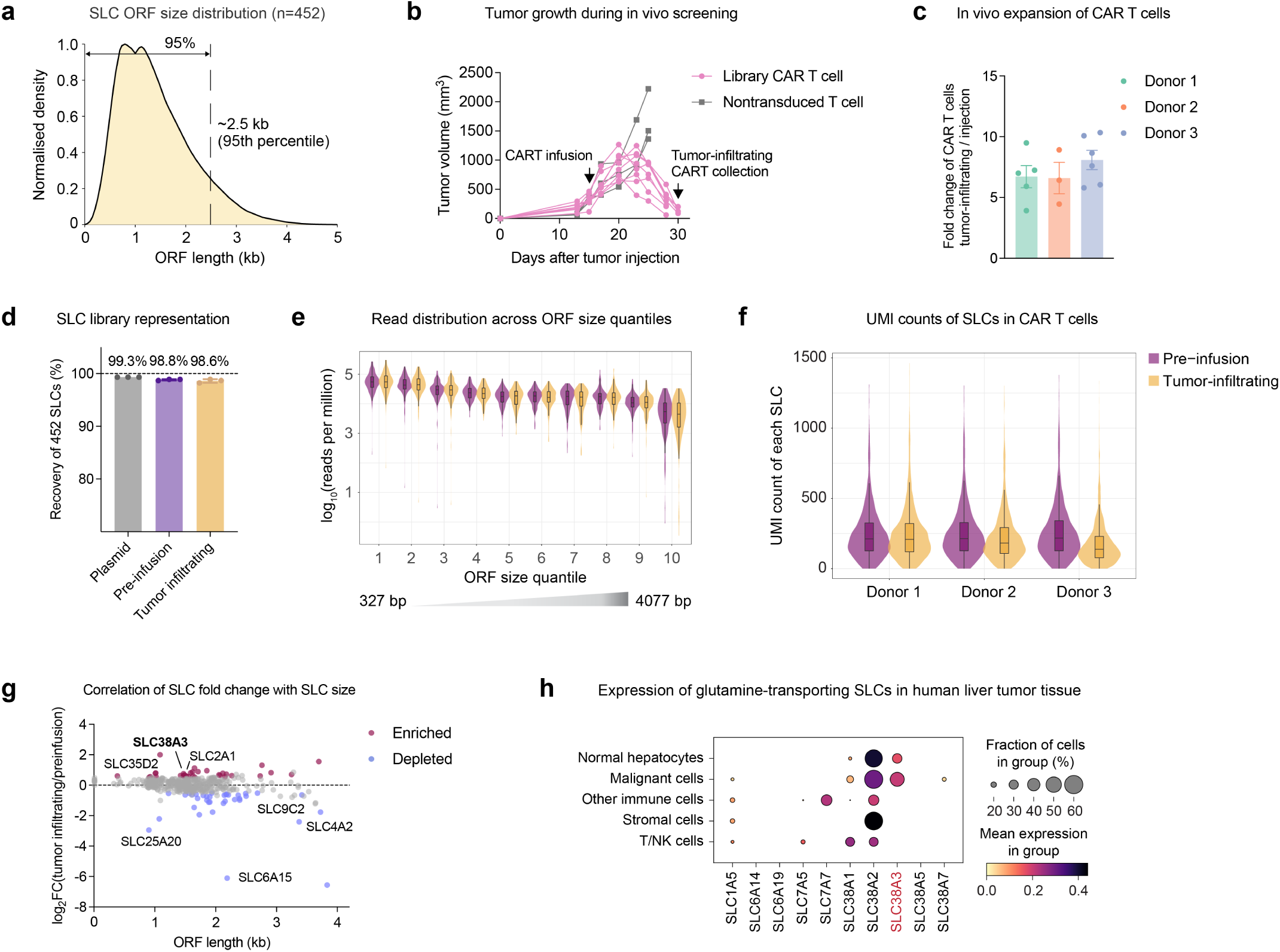
Ǫuality control metrics support the validity of the in vivo screening. (a) Histogram showing the distribution of ORF sizes in the SLC library. The majority of SLC ORFs fall between 0.9-1.5 kb, with 95% of all ORFs being smaller than 2.5 kb. (b) Tumor growth curves for tumor-bearing mice in response to treatment with the CAR-SLC T cells of the in vivo screening. The CAR-SLC T cells were injected 2 weeks after tumor cell injection, and tumor-infiltrating CAR T cells were collected 2 weeks after treatment. Each line represents one mouse. (c) Fold change of the number of tumor-infiltrating CAR T cells versus the number of injected CAR T cells. Each dot represents one tumor sample. Data shown are for three independent T cell donors with three to six mice each. (d) Recovery of the 452 SLCs from the plasmid library, pre-infusion CAR T cells, and tumor-infiltrating CAR T cells. Each dot represents the average for one donor (three T cell donors with at least three replicates each). (e) Violin plots showing the distribution of barcode sequencing reads for ORFs of different sizes. ORFs were assigned to ten size quantiles, from smallest (327 bp) to largest (4077 bp). Purple indicates pre-infusion CAR T cells and orange indicates tumor-infiltrating CAR T cells (three T cell donors with at least three replicates each). (f) Violin plots showing the distribution of UMI counts per SLC for pre-infusion and tumor-infiltrating CAR T cells. Purple, pre-infusion CAR T cells; orange, tumor-infiltrating CAR T cells. Each donor had at least three replicates. (g) Scatter plot illustrating the absence of a strong association between SLC abundance change and ORF size. Purple dots indicate significantly enriched SLCs (log_2_FC > 0.5, FDR-adjusted *p*-value < 0.05), blue dots indicate significantly depleted SLCs (log_2_FC < -0.5, FDR-adjusted *p*-value < 0.05), and grey dots indicate unchanged SLCs. Screens were performed independently for three T cell donors with three to six mice each. (h) Expression of SLC genes related to glutamine metabolism in the indicated cell populations from human liver tumor tissue, based on published single-cell RNA-seq data (GSE149614). Error bars in (c) and (d) show mean ± SEM.

**Figure S2.**
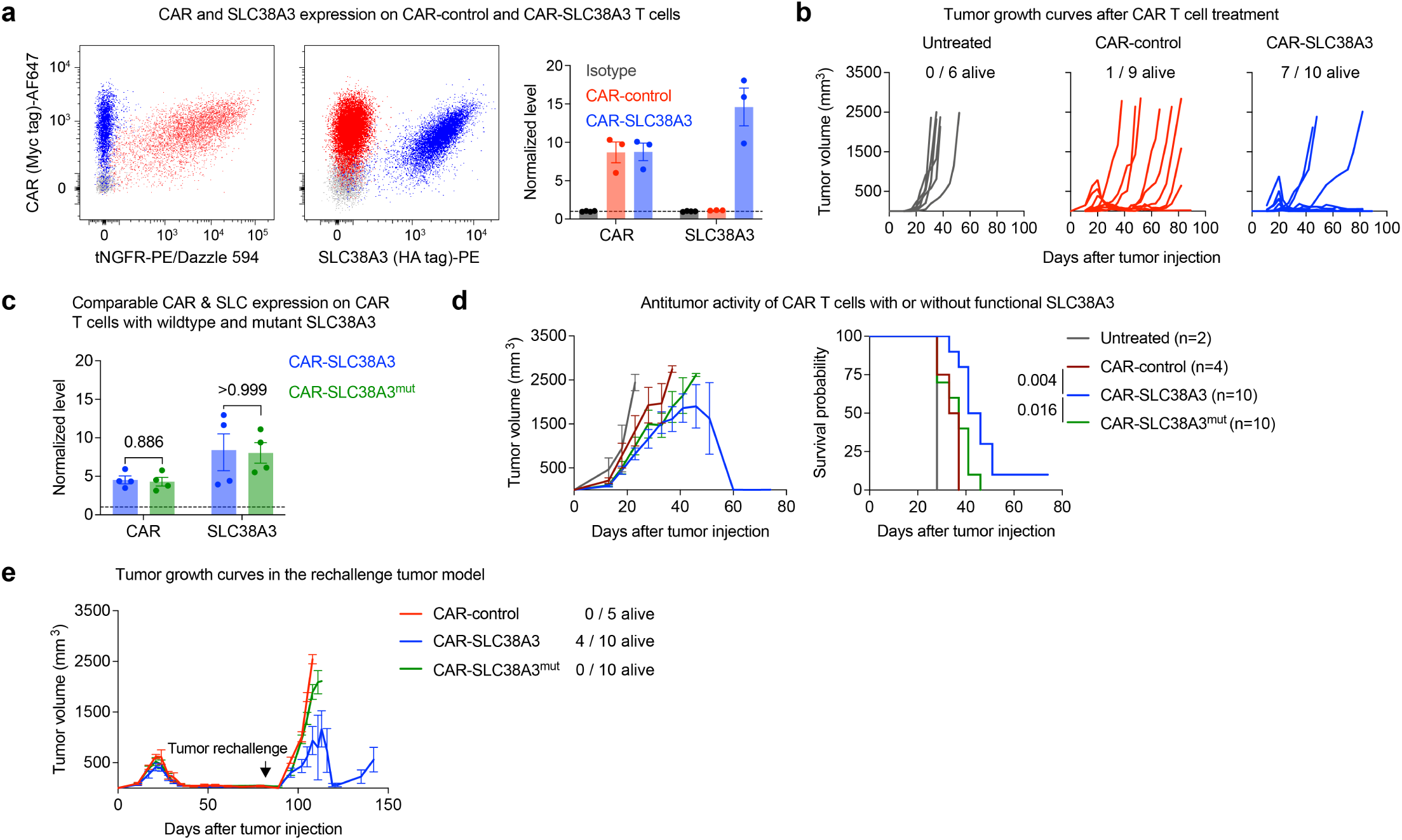
SLC38A3 expression promotes long-term antitumor efficacy of CAR T cells. (a) Representative flow cytometry plots (left) and bar plots (right) showing the expression of the CAR and of either SLC38A3 (*CAR-SLC38A3*, blue) or tNGFR used as control (*CAR-control*, red), following their co-delivery with the CARGO vector (n = 3 across three T cell donors). (b) Tumor growth curves for CAR T cell treatment of xenograft liver tumors (Huh7) as illustrated in Fig. 1d. Experiments were performed independently for two T cell donors with five mice each. Text labels show the number of surviving mice at the end of the experiment versus the total number of mice. Each line represents one mouse. (c) CAR and SLC38A3 expression on CAR T cells expressing either wildtype SLC38A3 (blue) or the transport-deficient SLC38A3^mut^ variant (green) (n = 4 across two T cell donors). (d) Tumor growth curves (left) and survival curves (right) for CAR T cell treatment of xenograft liver tumors (Huh7). Two million tumor cells were injected subcutaneously into mice to induce high-burden tumor formation. Established tumors were treated two weeks later with 0.5 million CAR T cells injected into the tail vein. Experiments were performed independently for two T cell donors with five mice each. (e) Tumor growth curves for the tumor rechallenge model depicted in Fig. 1f. The black arrow indicates when Huh7 cells were injected into the opposite flank (day 84). Experiments were performed independently for two T cell donors. Text labels show the number of surviving mice at the end of the experiment versus their total number. *P*-values in (c) were determined by two-tailed Mann-Whitney tests. *P*-values in (d) were determined by log-rank (Mantel-Cox) tests. Error bars in (a), (c), (d), and (e) show mean ± SEM.

**Figure S3.**
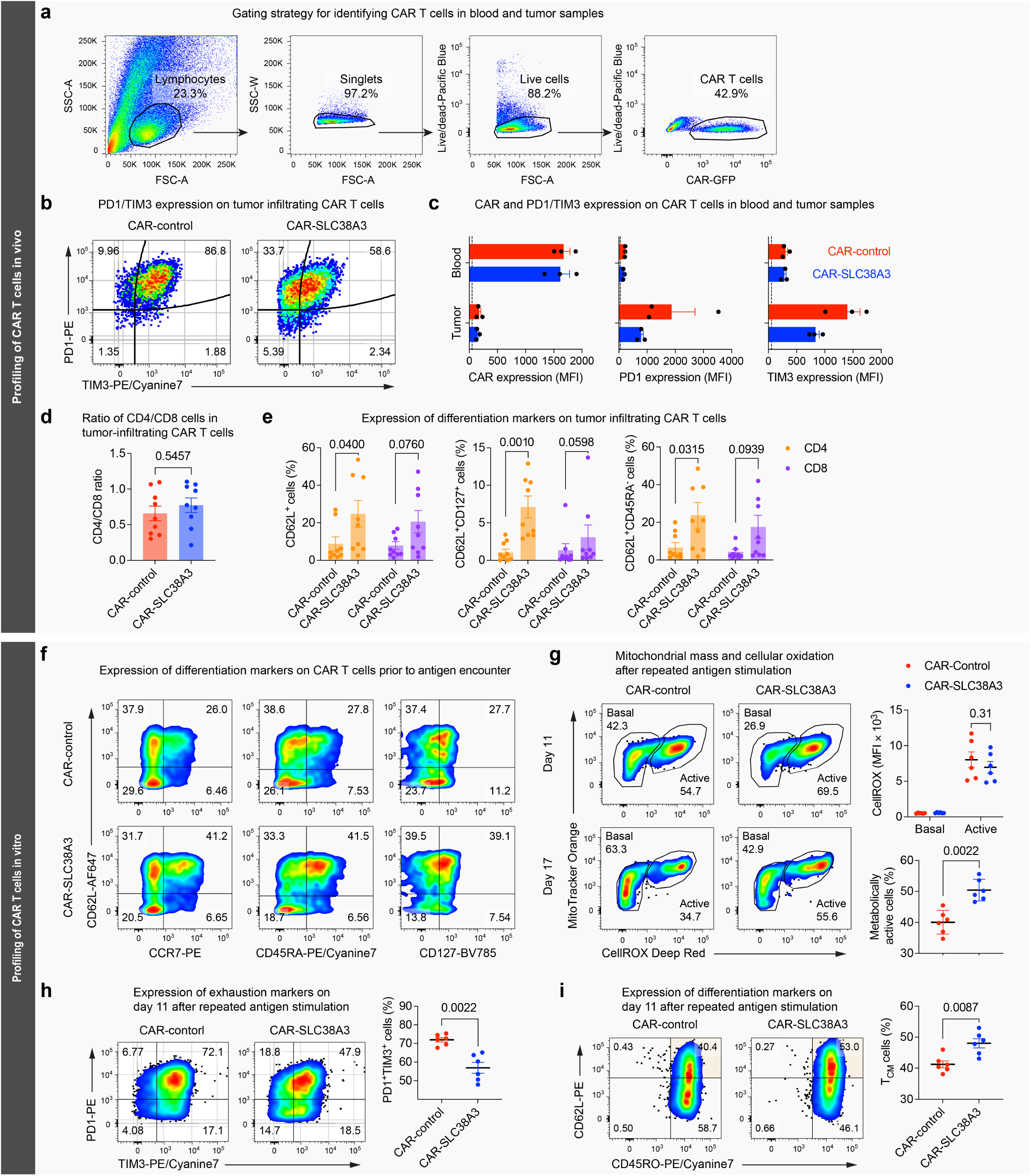
SLC38A3 expression promotes memory cell differentiation and metabolic fitness while mitigating exhaustion in CAR T cells. (a) Gating strategy for identifying CAR T cells in blood and tumor samples. Cells were selected based on forward scatter (FSC-A) and side scatter (SSC-A), cell doublets were excluded, and viable cells were gated based on live/dead staining with DAPI (Pacific Blue). CAR T cells were then identified based on their GFP expression. (b) Flow cytometry plots of PD-1 and TIM-3 expression on tumor-infiltrating CAR T cells. Representative data from three independent experiments are shown. (c) Bar plots of CAR, PD-1, and TIM-3 cell-surface expression levels on CAR T cells isolated from blood and tumor samples. Each dot represents one mouse. (d) Bar plot showing the ratio of CD4^+^ to CD8^+^ cells among tumor-infiltrating CAR T cells. Each dot represents one mouse (n = 9 across three T cell donors). (e) Percentage of tumor-infiltrating CD4^+^ and CD8^+^ CAR T cells that are positive for cell-surface markers associated with central memory cell differentiation. Each dot represents one mouse (n = 9 across three T cell donors). (f) Flow cytometry plots of stem cell-like memory (SCM) marker expression on CAR T cells prior to tumor stimulation (in vitro). Representative data from at least three independent experiments are shown. (g) Metabolic state of CAR T cells after repeated antigen stimulation. Left, representative flow cytometry plots of CAR T cells stained with MitoTracker (mitochondrial mass) and CellROX (cellular oxidation). Basal: cells with low mitochondrial mass and oxidation; Active: cells with elevated mitochondrial mass and cellular oxidation. Upper right, quantification of CellROX signal in basal and active cell populations. Bottom right, percentage of metabolically active cells among CAR T cells on day 17 (n = 6 across three T cell donors). (h) Exhaustion marker expression on CAR T cells after repeated antigen stimulation (day 11). Left, representative flow cytometry plots of PD-1 and TIM-3 expression on CAR T cells. Right, percentage of PD-1^+^TIM-3^+^ cells among all CAR T cells (also shown in Fig. 2i) (n = 6 across three T cell donors). (i) Differentiation marker expression on CAR T cells after repeated antigen stimulation (day 11). Left, representative flow cytometry plots of CD62L and CD45RO expression on CAR T cells. Right, percentage of CD62L^+^CD45RO^+^ central memory (CM) cells among all CAR T cells (n = 6 across three T cell donors). *P*-values in this figure were determined by two-tailed Mann-Whitney tests. Error bars show mean ± SEM.

**Figure S4.**
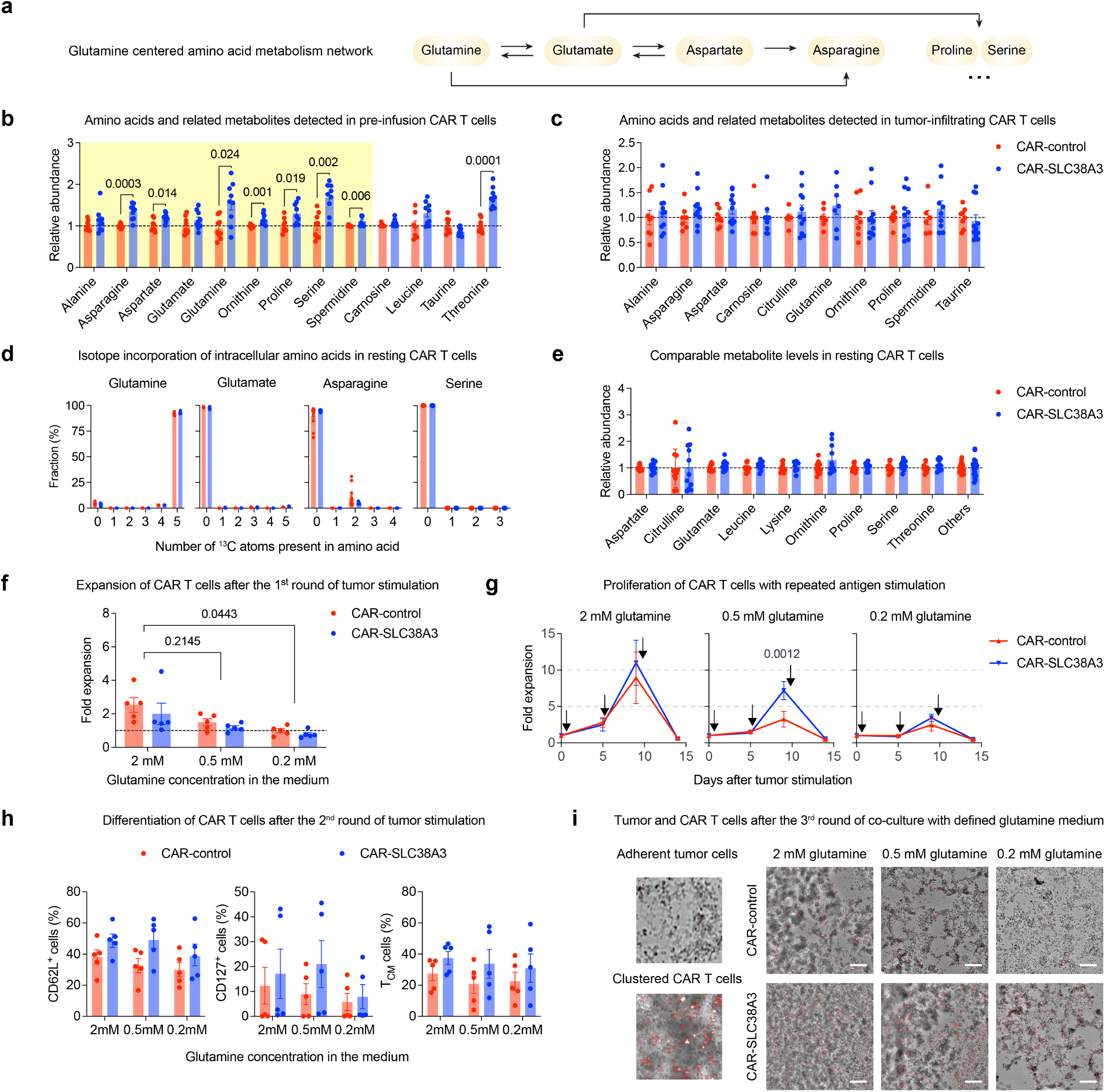
SLC38A3 expression modulates amino acid metabolism in CAR T cells and supports antitumor efficacy under glutamine restriction. (a) Schematic of glutamine-related amino acid metabolism. (b) Bar plots of intracellular amino acids and related metabolites in pre-infusion CAR T cells. The yellow area highlights amino acids involved in glutamine metabolism (n = 9 across three T cell donors). (c) Bar plots of intracellular amino acids and related metabolites in tumor-infiltrating CAR T cells (providing details for the “Others” group in Fig. 3b). Each dot represents one mouse (n = 11 across four T cell donors with two to three mice each). Missing dots are due to undetected signal. (d) Bar plots of ^13^C incorporation for glutamine metabolism-related amino acids (n = 12 across four T cell donors). (e) Bar plots of intracellular metabolites in resting CAR T cells (n = 12 across four T cell donors). (f) Bar plots of CAR T cell proliferation after the initial tumor stimulation under defined glutamine conditions (n = 5 across three T cell donors). (g) Line plots of CAR T cell proliferation after repeated stimulation under defined glutamine conditions (n = 5 across three T cell donors). Black arrows indicate the starting time of each stimulation round on days 0, 5, and 9. (h) Differentiation states among CAR T cells after the second round of tumor stimulation (n = 5 across three T cell donors). Central memory T cells were defined as CD62L^+^CD127^+^ cells. Two-way ANOVA found a significant difference for CD62L^+^ cells (*p*-value: 0.012), but no significant difference for glutamine concentration. (i) Representative images showing co-cultured CAR T cells (red) and tumor cells after the third round of tumor stimulation under defined glutamine conditions (as shown in Fig. 3g). Zoom-in images (left) with examples of adherent tumor cells and clustered CAR T cells from the broader image (right). Scale bars, 1 mm. *P*-values in (b) were determined by two-tailed Mann-Whitney tests, and *P*-values in (f), (g), and (h) were determined by two-way ANOVA tests with multiple-testing correction. Error bars show mean ± SEM.

**Figure S5.**
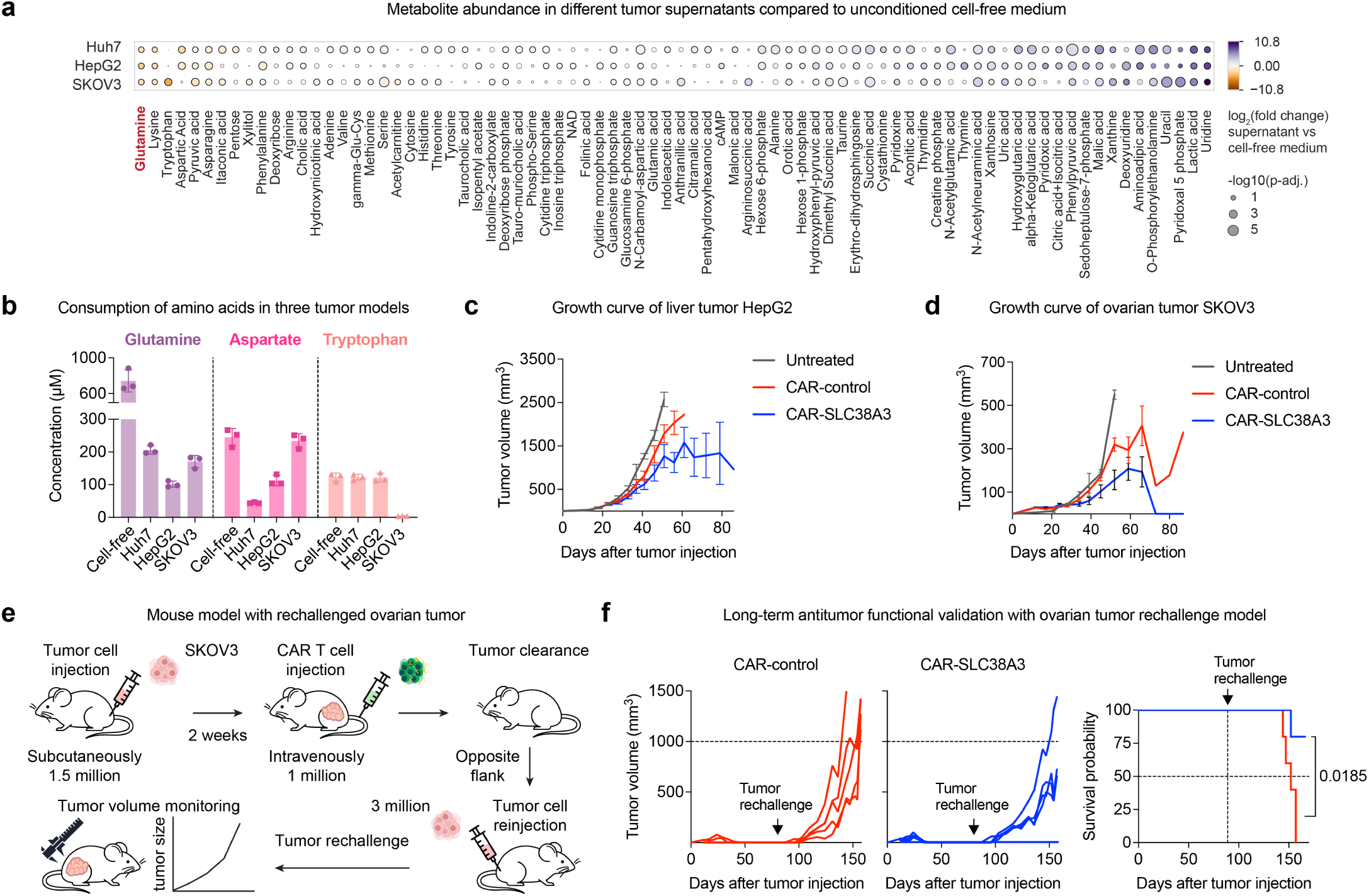
Validation experiments confirm the in vivo performance of SLC38A3-expressing CAR T cells for several tumor models with high glutamine consumption. (a) Heatmap showing metabolite changes for tumor cell-conditioned medium compared to unconditioned cell-free medium (n = 3, only metabolites with a significant change in at least one tumor cell type are shown, full results are in **Table S6**). (b) Bar plots of glutamine, aspartate, and tryptophan levels in unconditioned cell-free and tumor cell-conditioned medium (n = 3). (c) Tumor growth curves for CAR T cell treatment of HepG2 liver tumors as illustrated in Fig. 4c. Experiments were repeated independently across two T cell donors. Untreated group, n = 3; CAR T cell treated group, n = 10. (d) Tumor growth curves for CAR T cell treatment of SKOV3 ovarian tumors as illustrated in Fig. 4e. Experiments were repeated independently across four T cell donors. Untreated group, n = 3; CAR T cell treated group, n = 19. (e) Outline of the ovarian tumor rechallenge model for assessing persistence and long-term function of the CAR T cells. Mice that had cleared the initial tumor were rechallenged on day 90 by injecting SKOV3 tumor cells in the opposite flank without giving a new dose of CAR T cells, thus relying on persistent CAR T cells from the initial dose. (f) Tumor growth curves (left) and survival curves (right) for the ovarian tumor rechallenge model, showing mice previously injected with either control CAR T cells (red) or CAR T cells expressing SLC38A3 (blue). Each line (left) represents one mouse (n = 5). *P*-values in (a) and (f) were determined by t-tests with multiple-testing correction and separately by log-rank (Mantel-Cox) tests. Error bars in (b), (c), and (d) show mean ± SD.

## Supplementary Tables

**Table S1. Details of the SLC ORF library including gene names, barcodes, and sequences**

**Table S2. Barcode counts and differential analysis for the in vivo screening**

**Table S3. Unique molecular identifier (UMI) counts for the in vivo screening**

**Table S4. Metabolic profiling data for pre-infusion and tumor-infiltrating CAR T cells**

**Table S5. Metabolic profiling data for ^13^C-glutamine tracing assay in resting CAR T cells**

**Table S6. Metabolic profiling data for the tumor cell-conditioned media**

**Table S7. Primer sequences for library cloning and barcode sequencing**

**Table S8. Flow cytometry antibodies for cell surface phenotyping and intracellular staining**

**Table N1. Barcode counts and differential analysis for the in vitro screening**

## References

1. Cappell, K.M. & Kochenderfer, J.N. Long-term outcomes following CAR T cell therapy: what we know so far. Nat Rev Clin Oncol 20, 359–371 (2023).

2. June, C.H. & Sadelain, M. Chimeric Antigen Receptor Therapy. N Engl J Med 379, 64–73 (2018).

3. Albelda, S.M. CAR T cell therapy for patients with solid tumours: key lessons to learn and unlearn. Nat Rev Clin Oncol 21, 47–66 (2024).

4. Uslu, U. & June, C.H. Beyond the blood: expanding CAR T cell therapy to solid tumors. Nat Biotechnol 43, 506–515 (2025).

5. Peng, J.J., Wang, L.M., Li, Z.Y., Ku, C.L. & Ho, P.C. Metabolic challenges and interventions in CAR T cell therapy. Science Immunology 8 (2023).

6. Leone, R.D. & Powell, J.D. Metabolism of immune cells in cancer. Nat Rev Cancer 20, 516–531 (2020).

7. DePeaux, K. & Delgoffe, G.M. Metabolic barriers to cancer immunotherapy. Nat Rev Immunol 21, 785–797 (2021).

8. Pallett, L.J., Dimeloe, S., Sinclair, L.V., Byrne, A.J. & Schurich, A. A glutamine ‘tug-of-war’: targets to manipulate glutamine metabolism for cancer immunotherapy. Immunother Adv 1, ltab010 (2021).

9. Lim, A.R., Rathmell, W.K. & Rathmell, J.C. The tumor microenvironment as a metabolic barrier to effector T cells and immunotherapy. Elife 9 (2020).

10. Park, J., Hsueh, P.C., Li, Z.Y. & Ho, P.C. Microenvironment-driven metabolic adaptations guiding CD8+T cell anti-tumor immunity. Immunity 56, 32–42 (2023).

11. Kishton, R.J., Sukumar, M. & Restifo, N.P. Metabolic Regulation of T Cell Longevity and Function in Tumor Immunotherapy. Cell Metab 26, 94–109 (2017).

12. Mcphedran, S.J., Carleton, G.A. & Lum, J.J. Metabolic engineering for optimized CAR-T cell therapy. Nature Metabolism 6, 396–408 (2024).

13. Wang, M.R., Mu, W., Zhen, A. & Kitchen, S.G. CRISPR/Cas strategies to enhance CAR T-cell function and persistence via metabolic reprogramming. Trends Biotechnol 44, 2265–2278 (2026).

14. Giuffrida, L. et al. CRISPR/Cas9 mediated deletion of the adenosine A2A receptor enhances CAR T cell efficacy. Nat Commun 12, 3236 (2021).

15. Wu, M.H. et al. Deleting the mitochondrial respiration negative regulator MCJ enhances the efficacy of CD8(+) T cell adoptive therapies in pre-clinical studies. Nat Commun 15, 4444 (2024).

16. Ye, L. et al. A genome-scale gain-of-function CRISPR screen in CD8 T cells identifies proline metabolism as a means to enhance CAR-T therapy. Cell Metab 34, 595–614 e14 (2022).

17. Cesar-Razquin, A. et al. A Call for Systematic Research on Solute Carriers. Cell 162, 478–87 (2015).

18. Dobson, P.D. & Kell, D.B. Carrier-mediated cellular uptake of pharmaceutical drugs: an exception or the rule? Nat Rev Drug Discov 7, 205–20 (2008).

19. Pizzagalli, M.D., Bensimon, A. & Superti-Furga, G. A guide to plasma membrane solute carrier proteins. FEBS J 288, 2784–2835 (2021).

20. Bian, Y. et al. Cancer SLC43A2 alters T cell methionine metabolism and histone methylation. Nature 585, 277–282 (2020).

21. Cao, T.Y. et al. Cancer SLC6A6-mediated taurine uptake transactivates immune checkpoint genes and induces exhaustion in CD8+T cells. Cell 187, 2288–2304 (2024).

22. Han, C. et al. Cystine deprivation triggers CD36-mediated ferroptosis and dysfunction of tumor infiltrating CD8(+) T cells. Cell Death Dis 15, 145 (2024).

23. Shi, Y.Z. et al. GLUT1 overexpression enhances CAR T cell metabolic fitness and anti-tumor efficacy. Molecular Therapy 32, 2393–2405 (2024).

24. Guerrero, J.A. et al. GLUT1 overexpression in CAR-T cells induces metabolic reprogramming and enhances potency. Nature Communications 15, 8658 (2024).

25. Panetti, S. et al. Engineering amino acid uptake or catabolism promotes CAR T-cell adaption to the tumor environment. Blood Adv 7, 1754–1761 (2023).

26. Wolf, G. et al. The efflux pump ABCC1/MRP1 constitutively restricts PROTAC sensitivity in cancer cells. Cell Chemical Biology 32, 291–306 (2025).

27. Joung, J. et al. A transcription factor atlas of directed differentiation. Cell 186, 209–229 e26 (2023).

28. Konermann, S. et al. Genome-scale transcriptional activation by an engineered CRISPR-Cas9 complex. Nature 517, 583–8 (2015).

29. Sanson, K.R. et al. Optimized libraries for CRISPR-Cas9 genetic screens with multiple modalities. Nat Commun 9, 5416 (2018).

30. Knudsen, N.H. et al. In vivo CRISPR screens identify modifiers of CAR T cell function in myeloma. Nature 646, 953–962 (2025).

31. Datlinger, P. et al. Systematic discovery of CRISPR-boosted CAR T cell immunotherapies. Nature 646, 963–972 (2025).

32. Parks, S.K., Chiche, J. & Pouyssegur, J. Disrupting proton dynamics and energy metabolism for cancer therapy. Nat Rev Cancer 13, 611–23 (2013).

33. Hadley, B. et al. Nucleotide Sugar Transporter SLC35 Family Structure and Function. Computational and Structural Biotechnology Journal 17, 1123–1134 (2019).

34. Gu, S., Roderick, H.L., Camacho, P. & Jiang, J.X. Identification and characterization of an amino acid transporter expressed differentially in liver. Proc Natl Acad Sci U S A 87, 3230–5 (2000).

35. Fei, Y.J. et al. Primary structure, genomic organization, and functional and electrogenic characteristics of human system N 1, a Na+- and H+-coupled glutamine transporter. J Biol Chem 275, 23707–17 (2000).

36. Chaudhry, F.A. et al. Molecular analysis of system N suggests novel physiological roles in nitrogen metabolism and synaptic transmission. Cell 99, 769–80 (1999).

37. Lu, Y. et al. A single-cell atlas of the multicellular ecosystem of primary and metastatic hepatocellular carcinoma. Nat Commun 13, 4594 (2022).

38. Chan, K. et al. Loss of function mutation of the Slc38a3 glutamine transporter reveals its critical role for amino acid metabolism in the liver, brain, and kidney. Pflugers Archiv-European Journal of Physiology 468, 213–227 (2016).

39. Fraietta, J.A. et al. Determinants of response and resistance to CD19 chimeric antigen receptor (CAR) T cell therapy of chronic lymphocytic leukemia. Nat Med 24, 563–571 (2018).

40. Xu, Y. et al. Closely related T-memory stem cells correlate with in vivo expansion of CAR.CD19-T cells and are preserved by IL-7 and IL-15. Blood 123, 3750–9 (2014).

41. Biasco, L. et al. Clonal expansion of T memory stem cells determines early anti-leukemic responses and long-term CAR T cell persistence in patients. Nat Cancer 2, 629–642 (2021).

42. Rubio-Aliaga, I. & Wagner, C.A. Regulation and function of the SLC38A3/SNAT3 glutamine transporter. Channels 10, 440–452 (2016).

43. Ananieva, E.A., Powell, J.D. & Hutson, S.M. Leucine Metabolism in T Cell Activation: mTOR Signaling and Beyond. Adv Nutr 7, 798S–805S (2016).

44. Nicklin, P. et al. Bidirectional transport of amino acids regulates mTOR and autophagy. Cell 136, 521–34 (2009).

45. Cluntun, A.A., Lukey, M.J., Cerione, R.A. & Locasale, J.W. Glutamine Metabolism in Cancer: Understanding the Heterogeneity. Trends Cancer 3, 169–180 (2017).

46. Altman, B.J., Stine, Z.E. & Dang, C.V. From Krebs to clinic: glutamine metabolism to cancer therapy. Nat Rev Cancer 16, 619–634 (2016).

47. Yang, L.F., Venneti, S. & Nagrath, D. Glutaminolysis: A Hallmark of Cancer Metabolism. Annual Review of Biomedical Engineering 19, 163–194 (2017).

48. Newick, K., O’Brien, S., Moon, E. & Albelda, S.M. CAR T Cell Therapy for Solid Tumors. Annu Rev Med 68, 139–152 (2017).

49. Bader, J.E., Voss, K. & Rathmell, J.C. Targeting Metabolism to Improve the Tumor Microenvironment for Cancer Immunotherapy. Mol Cell 78, 1019–1033 (2020).

50. Hanahan, D. Hallmarks of Cancer: New Dimensions. Cancer Discov 12, 31–46 (2022).

51. Pavlova, N.N. & Thompson, C.B. The Emerging Hallmarks of Cancer Metabolism. Cell Metab 23, 27–47 (2016).

52. DeBerardinis, R.J. & Chandel, N.S. Fundamentals of cancer metabolism. Sci Adv 2, e1600200 (2016).

53. Boulland, J.L. et al. Cell-specific expression of the glutamine transporter SN1 suggests differences in dependence on the glutamine cycle. Eur J Neurosci 15, 1615–31 (2002).

54. Uhlen, M. et al. Proteomics. Tissue-based map of the human proteome. Science 347, 1260419 (2015).

55. Johnson, M.O. et al. Distinct Regulation of Th17 and Th1 Cell Differentiation by Glutaminase-Dependent Metabolism. Cell 175, 1780–1795 e19 (2018).

56. Wang, R. et al. The transcription factor Myc controls metabolic reprogramming upon T lymphocyte activation. Immunity 35, 871–82 (2011).

57. Ma, E.H. et al. (13)C metabolite tracing reveals glutamine and acetate as critical in vivo fuels for CD8 T cells. Sci Adv 10, eadj1431 (2024).

58. Wise, D.R. & Thompson, C.B. Glutamine addiction: a new therapeutic target in cancer. Trends Biochem Sci 35, 427–33 (2010).

59. Hushmandi, K. et al. Amino acid transporters within the solute carrier superfamily: Underappreciated proteins and novel opportunities for cancer therapy. Molecular Metabolism 84 (2024).

60. Jin, J., Byun, J.K., Choi, Y.K. & Park, K.G. Targeting glutamine metabolism as a therapeutic strategy for cancer. Experimental and Molecular Medicine 55, 706–715 (2023).

61. Wenes, M. et al. The mitochondrial pyruvate carrier regulates memory T cell differentiation and antitumor function. Cell Metabolism 34, 731–746 (2022).

62. Demicco, M., Liu, X.Z., Leithner, K. & Fendt, S.M. Metabolic heterogeneity in cancer. Nat Metab 6, 18–38 (2024).

63. Faubert, B., Solmonson, A. & DeBerardinis, R.J. Metabolic reprogramming and cancer progression. Science 368, eaaw5473 (2020).

64. Amendola, M., Venneri, M.A., Biffi, A., Vigna, E. & Naldini, L. Coordinate dual-gene transgenesis by lentiviral vectors carrying synthetic bidirectional promoters. Nature Biotechnology 23, 108–116 (2005).

65. Zhu, A.X. et al. First-in-Man Phase I Study of GC33, a Novel Recombinant Humanized Antibody Against Glypican-3, in Patients with Advanced Hepatocellular Carcinoma. Clinical Cancer Research 19, 920–928 (2013).

66. Albanell, J. & Baselga, J. Trastuzumab, a humanized anti-HER2 monoclonal antibody, for the treatment of breast cancer. Drugs Today (Barc) 35, 931–46 (1999).

67. Lynn, R.C. et al. c-Jun overexpression in CAR T cells induces exhaustion resistance. Nature 576, 293–300 (2019).

68. Doan, A.E. et al. FOXO1 is a master regulator of memory programming in CAR T cells. Nature 629, 211–218 (2024).

69. Zhao, Z.G. et al. Structural Design of Engineered Costimulation Determines Tumor Rejection Kinetics and Persistence of CAR T Cells. Cancer Cell 28, 415–428 (2015).

70. Bueschl, C. et al. PeakBot: machine-learning-based chromatographic peak picking. Bioinformatics 38, 3422–3428 (2022).

71. Wolf, F.A., Angerer, P. & Theis, F.J. SCANPY: large-scale single-cell gene expression data analysis. Genome Biology 19 (2018).

72. Fernandez, C.A., Des Rosiers, C., Previs, S.F., David, F. & Brunengraber, H. Correction of 13C mass isotopomer distributions for natural stable isotope abundance. J Mass Spectrom 31, 255–62 (1996).

73. Buescher, J.M. et al. A roadmap for interpreting (13)C metabolite labeling patterns from cells. Curr Opin Biotechnol 34, 189–201 (2015).

