## Supplementary Note for "In vivo screening identifies SLC38A3 as a metabolic booster for CAR T cell therapy"

### Supplementary Note. Hits from SLC in vitro screening enhance CAR T cell function in vitro but not in vivo

To discover metabolic boosters of CAR T cell function, we performed in vitro screening using a repeated antigen stimulation assay that seeks to mimic chronic tumor stimulation (**Fig. N1a**), using the same CAR T cells, SLC ORF library, and tumor cells (Huh7) as for the in vivo screening (**Fig. 1**). We identified multiple SLC transmembrane transporters that were significantly enriched or depleted in this screen (**Fig. N1b,c and Table N1**). Most notably, we observed strong enrichment of SLC39 family members, which control cellular zinc homeostasis[1], and of mitochondrial pyruvate carrier (MPC) genes, which modulate pyruvate entry into mitochondria[2].

In vitro validation experiments confirmed that MPC overexpression enhances CAR T cell expansion, albeit not as strongly as overexpression of the established and effective CAR T cell booster c-Jun[3] (**Fig. N1d**). However, this in vitro benefit did not translate to our subsequent in vivo validation experiments. Overexpression of MPC1 or MPC2, separately or in combination, failed to enhance antitumor function relative to control CAR T cells in Huh7 tumor-bearing mice (**Fig. N1e,f**). This discordance between in vitro expansion and in vivo efficacy underscores the limitations of cell culture models and highlights the relevance of in vivo screening to identify metabolic modulators that are effective within the in vivo tumor microenvironment.

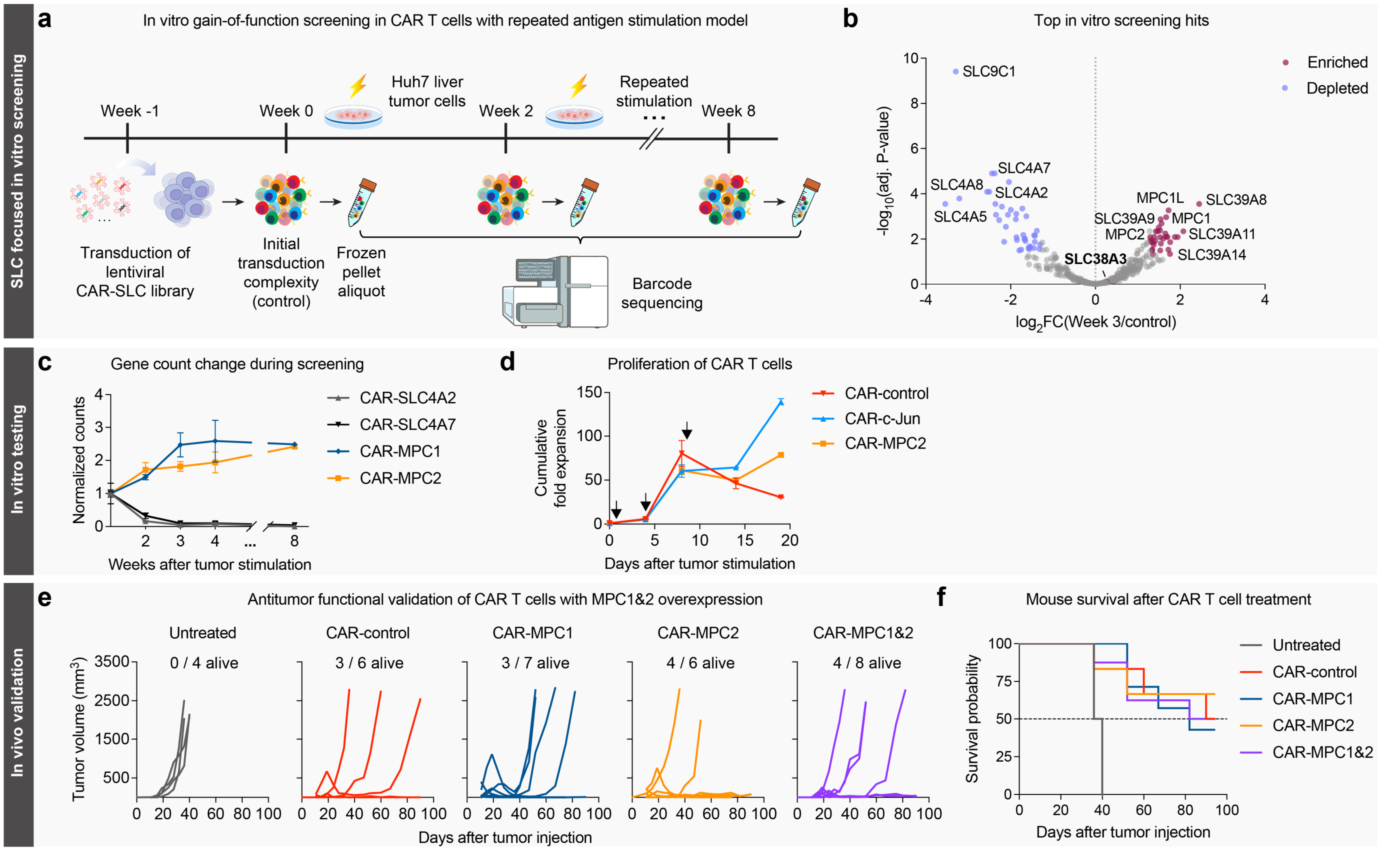


**Figure N1. In vitro screening identifies SLCs that enhance CAR T cell performance in vitro but not in vivo**

(a) Outline of in vitro gain-of-function screening in CAR T cells using a repeated antigen stimulation model.

(b) Volcano plot depicting changes in SLC representation in CAR T cells at week 3 compared to CAR T cells at baseline (week 0 control). Red points indicate significantly enriched SLCs ($log_2FC > 1.3$, FDR-adjusted $p$-value $< 0.05$); blue points indicate significantly depleted SLCs ($log_2FC < -1.3$, FDR-adjusted $p$-value $< 0.05$); gray points indicate unchanged SLCs. Screens were performed for three T cell donors with three replicates each.

(c) Temporal changes in normalized barcode counts in CAR T cells during in vitro screening (three human T cell donors with three technical replicates each).

(d) Cumulative fold expansion of CAR T cells following repeated antigen stimulation every 4 days. Black arrows indicate stimulations with Huh7 cells at an effector-to-target (E:T) ratio of 1:3 (three human T cell donors).

(e) Tumor growth curves for CAR T cell treatment of xenograft liver tumors (Huh7). 1 million Huh7 cells were injected subcutaneously into mice to induce tumor formation. Established tumors were treated two weeks later with 1 million CAR T cells injected into the tail vein. Each line represents one mouse. Experiments were performed independently for two T cell donors. The numbers indicate the number of surviving mice out of the total mice per group at the end of the experiment.

(f) Survival curves for the mice treated with CAR T cells overexpressing the indicated genes, as in (e). No statistically significant differences were detected between groups ($p$-value > 0.05, log-rank test).

Error bars in (c) and (d) show mean ± SD.

**References**


1. Jeong, J., and Eide, D.J. (2013). The SLC39 family of zinc transporters. *Mol Aspects Med 34*, 612-619.
2. Lacabanne, D., Ruprecht, J.J., Sichrovsky, M., Forrest, L.R., Leone, V., Tavoulari, S., and Kunji, E.R.S. (2026). Structural transport and inhibition mechanism of the mitochondrial pyruvate carrier. *Trends Biochem Sci 51*, 142-157.
3. Lynn, R.C., Weber, E.W., Sotillo, E., Gennert, D., Xu, P., Good, Z., Anbunathan, H., Lattin, J., Jones, R., Tieu, V., et al. (2019). c-Jun overexpression in CAR T cells induces exhaustion resistance. *Nature 576*, 293-300.
